# Male obesity causes adipose mitochondrial dysfunction in F_1_ progeny via a let-7-DICER axis

**DOI:** 10.1101/2024.10.03.615866

**Authors:** Chien Huang, Joo Hyun Park, Ali Altıntaş, Natasa Stanic, Kristine Kyle de Leon, Signe Isacson, Panagiotis Kalogeropoulos, Hande Topel Batarlar, Rocio Valdebenito Malmros, Jesper Havelund, Bjørk Ditlev Marcher Larsen, Yen-Ting Chien, Wen-Chi Huang, Karolina Szczepanowska, Jan-Wilm Lackmann, Aleksandra Trifunovic, Eva Kildall Hejbøl, Sönke Detlefsen, Ida Engberg Jepsen, Stefanie Hansborg Kolstrup, Ricardo Laguna-Barraza, Javier Martin-Gonzalez, Konstantin Khodosevich, Nils J. Færgeman, Marcelo A. Mori, Marc R. Friedländer, Anita Oest, Romain Barrès, Jan-Wilhelm Kornfeld

## Abstract

We here describe that obesity and weight loss in male mice cause reversible abnormalities in glucose and lipid metabolism, serum metabolomes and lipidomes as well as expression of microRNAs, mRNAs and proteins controlling mitochondrial function in epididymal white adipose tissue. When mating obese male mice with lean females, we observed reductions in expression and translation of genes encoding mitochondrial respiratory components in (F_1_) offspring that closely resemble those observed in the paternal (F_0_) generation. When mapping miRNA regulation across somatic organs (i.e., liver, adipose) and sperm and F_0/1_ generations, we found that obesity and weight loss reversibly affected miRNA levels, and that *let-7* isoforms were induced in obese F_0_ and F_1_ adipose tissue and sperm of obese F_0_ mice, eliciting qualitatively similar responses in two adjacent tissues. Overexpressing *let-7* in adipocytes silenced DICER1, a miRNA processing enzyme crucial for adipose adaptation to obesity as evidenced by deficiencies in mitochondrial function following DICER1 loss in primary adipocytes. Also, microinjection of synthetic *let-7* mimetics at physiological levels found in obese sperm into zygotes from lean mice elicited glucose intolerance and impediments in adipose mitochondrial gene expression in mice sired from *let-7* microinjected zygotes, phenocopying hereditary aspects of paternal obesity. When performing single-cell RNA-Seq of miRNA-injected embryos, *let-7* impaired mitochondrial gene expression, suggesting altered oxidative metabolism following zygotic *let-7* delivery. When studying miRNA alterations in human semen, lifestyle-induced weight loss downregulated *hsa-let-7*, suggesting similar roles for human *let-7* in gametic epigenomes and embryogenesis.

## Introduction

### Intergenerational epigenetic inheritance of metabolic dysfunction

Genome-wide association studies have demonstrated that genetic (i.e., DNA) variants only explain 20% of obesity heritability^1^. Given pandemic rises in obesity in evolutionarily relatively short timescales, it was proposed that non-genetic or ‘*epi-genetic*’ processes, that are spurred by obesogenic lifestyles and other changes in environment exposures, might increase obesity susceptibility and contribute to disease incidences. This form of epigenetic inheritance, if convincingly demonstrated^2^, is particularly intriguing for cases of paternal programming where affected fathers contribute only sperm as biological material to embryos (intergenerational inheritance) or no genetic/biological material at all (transgenerational inheritance). Epidemiological support for the existence of hereditary, paternally-acquired traits comes from historical records demonstrating that ample ancestral food supplies affects cardiometabolic traits up to two generations^3,4^, showing that individuals carry increased risks of cardiometabolic dysfunction if their parents or grandparents were obese or were malnourished^5^. In line with correlative observations in human, mechanistic studies in isogenic (genetically identical) mouse strains like C57BL76 demonstrated that paternal obesity can impair metabolism in offsprings (F_1_ generation and beyond) and, specific diet like low-protein^6^ or high-fat diet feeding^7–9^ and stress-related cues such as depression-like behavior^10^, fear-conditioning^11^ and psychological traumata^12^ impart specific epigenetic behavioral and metabolic effects in offsprings. Recently, beneficial types of epigenetic programming in adipose tissue were also demonstrated where offsprings of mice exposed to cold temperatures display increased amounts of metabolically favourable brown adipose tissue ^13^, suggesting an unexpectedly high plasticity of the molecular and cellular mechanisms underlying intergenerational epigenetic responses.

### The molecular basis for epigenetic inheritance of paternal obesity in male gametes

The molecular and cellular basis for epigenetic heredity remains largely enigmatic: Although changes in epigenetic marks like methylation of genomic DNA, histone post-translational modifications and alterations in small Noncoding RNAs (sncRNAs) correlate with hereditary processes in male gametes^14^. Although lifestyle/diet-induced changes to DNA methylation and histones modifications were reported, the evidence for causatively implicating these in eliciting paternal response in progeny is rare for higher mammalian species like rodents and humans. Other emerging instigators of intergenerational response are found amongst the different types of gametic sncRNAs such as microRNAs (miRNAs): miRNAs in gametes were initially identified and functionally dissected in nematodes and fruit flies^15–18^, thus species where sncRNAs-evoked gametic gene silencing processes are well-understood ^19^. In contrast to DNA-linked marks, sncRNAs are very mobile in nature and are subject to microvesicular trafficking between cells and organs and secreted by somatic cell types in testes^20,21^. Thus, due to their high abundance of miRNAs in sperm, their dynamic changes following diet exposures^22,23^, their exchange between sperm and oocytes^23^ sperm-born sncRNAs have the unique potential for exchanging information about past lifestyle choices to zygotes and F_1_ progeny. Consistent with this, metabolic trait/disease-associated miRNAs can affect intergenerational phenotypes and phenocopy F_0_ lifestyles if microinjected ^24,25^ or removed from zygotes^26^. Beyond controlling gametic epigenomes, miRNAs are also essential for germ cell development processes such as spermatogenesis^27^ and embryogenesis^28^, and miRNA are exchanged between zygotes and other cell types ^12,25,29–31^. Taken together, it is plausible to hypothesize that obesity-evoked changes in sperm miRNAs alter zygotic miRNA pools and embryonic development, instigating epigenetic phenotypic traits and somatic gene expression in offsprings. As diverse paternal lifestyle exposures ranging from inflammation^15^, nutrition^29,31^ to stress ^12,30^ alter gametic miRNA repertoires and F_1_ phenotypes in non-overlapping ways, the intriguing possibility exists that environmental experiences are encoded in individual gametic sncRNA signature and that these sperm sncRNAs have the potential to transmit complex lifestyle exposures given the high bandwidth of RNA-mediated effects^32^.

### Regulation of adipose tissue (mitochondrial) function by miRNAs

Adipose tissue is a multifunctional endocrine organ that coordinates tissue-level (autocrine, juxtacrine, paracrine) and systemic (endocrine) responses by release of adipokines, lipids and inflammatory factors and, shown recently, miRNA-containing microvesicles (exosomes)^33–35^. It is believed that adipose tissue can safely store ingested nutrients as lipids, although chronic calorie excess causes insulin resistance^36^, aberrant gene expression in adipocytes^37^, vascular remodelling and immune cell infiltration^33^. Mitochondria in adipose tissue exert important functions for tissue/energy homeostasis, particularly during nutrient deprivation/excess^35^: Whereas brown adipose tissue (BAT) contains large numbers of mitochondria to generate heat by dissipation of the electron transport chain (ETC) proton gradients^38^, inguinal (beige, iWAT) and epididymal (white, eWAT) adipose tissue protects from calorie overload by removing nutrients from circulation and storage as energetic, indolent macromolecules such as triglycerides. Mitochondrial function in WAT is impaired in obesity ^39^ and reduced mitochondrial function in fat causes metabolic imbalance^40–43^. Mitochondrial DNA content, oxidative phosphorylation (OxPhos), abundance and composition of ETC complexes are all functionally impaired in obesity and morphological aberrations in mitochondrial structures are found in obesity^39,43^ and aging^44^. Conversely, treatment with the AMP-activated protein kinase (AMPK) agonist Metformin improves mitochondrial function in adipose tissue, as do exercise or calorie restriction, and all these interventions correlate with metabolic improvement^45^. Recently, miRNAs were identified as important regulators of mitochondrial formation and function and known ‘mito-miRs’ like *miR-143, miR-181a, miR-378* instigate, whereas *miR-532* which impairs mitochondrial function by (post-transcriptionally) silencing mitochondrial translocation and mt-RNA homeostasis^46–48^. We and others showed that removing miRNAs from adipocytes by conditionally ablating DICER1, a central component of the conserved miRNA processing machinery, impairs adipogenesis, triggering lipodystrophy and metabolic dysfunction in mice^49–51^.

Collectively, we report that obesity in male mice is linked to reversible miRNA, mRNA and proteins alterations in adipose mitochondrial gene regulation that match those in adipose tissue of lean isogenic F_1_ progeny. Using integrative analyses of adipose, liver and sperm miRNA:mRNA gene networks, we identify *let-7* as induced by obesity across cell types and generations and mechanistically implicate *let-7* driven suppression of DICER1 and concomitant declines in mitochondrial function in adipocytes in inherited mouse phenotypes, thereby establishing sperm-born *let-7* as intergenerational driver of mitochondrial dysfunction and glucose impairments in F_1_ offspring.

## Results

### Paternal obesity elicits intergenerational impairments in adipose mitochondrial function

Whilst intergenerational effects of paternal (F_0_) obesity on offspring (F_1_) phenotypes and metabolic health have been reported, the specific organ-level manifestations of this dysfunction in progeny, and the precise somatic and germ cell-intrinsic molecular mechanisms in obese mice and their descendants, remain surprisingly elusive. To investigate this using isogenic rodents (*Mus musculus*), we used diet-induced obese (DIO) and weight-regressed C57BL/6N male mice, and fed 6-week-old animals with 1) low-fat diet (LF) for 18 weeks (F_0_ LF), 2) high-fat diet (HF) for 18 weeks (F_0_ HF) or 3) HF for 9 weeks, followed by 9 weeks of LF-driven weight loss (F_0_ HF-LF, **Fig. 1a**) to allow at least three cycles of spermatogenesis. For each F_0_ group, we mated eight randomly selected mice with unexposed lean virgin, female C57BL/6N mice, and exposed both male and female F_1_ progeny of each group, i.e., F_1_ (F_0_ LF, n=44 pups), F_1_ (F_0_ HF, n=52 pups) and F_1_ (F_0_ HF-LF, n=38 pups) mice to LF after weaning to test for metabolic and metabolic sequelæ of F_0_ diet exposure and removal. We did not additionally challenge F_1_ mice with HF given that effects of HF and epigenetic F_0_ effects were reported as cumulative with regards to F_1_ phenotypes ^52^ (**Fig. 1b**). Noteworthy, we detected no alterations in fecundity, male-to-female pup ratio and litters sizes (**Fig.S1a,b**), suggesting that F_0_ obesity did not affect development and embryonic implantation rates nor indirectly affected maternal provisioning after birth due to different litter sizes, two potential confounders.

**Figure 1:**
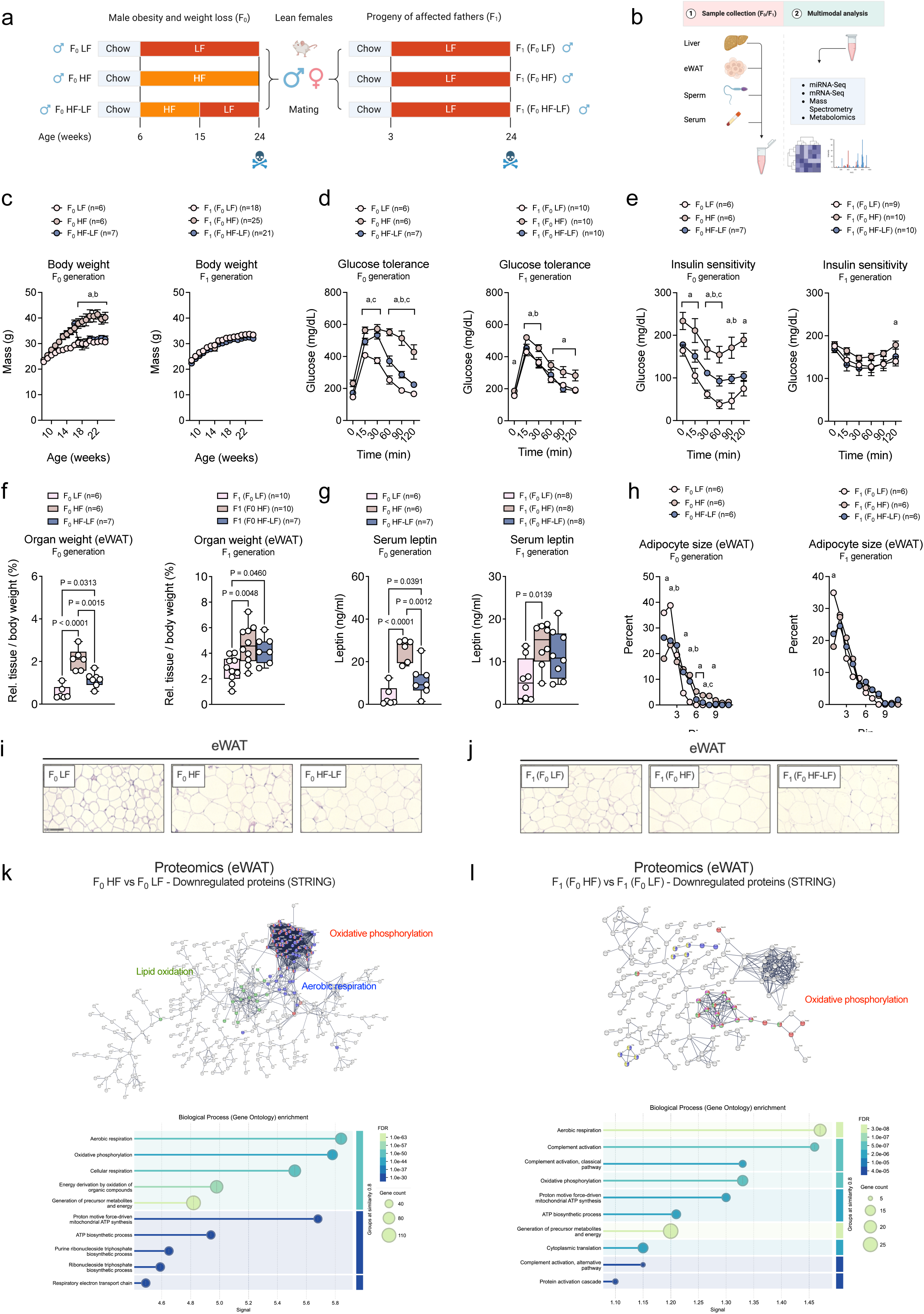
Paternal obesity elicits heritable impairments in glucose metabolism and adipose mitochondrial function. (**a,b**) Illustration of intergenerational *in vivo* cohorts depicting F_0_ groups of **(a)** paternal (F_0_) obesity and weight-regression, male and female F_1_ offspring mice **(b)** and analytical workflows incl. RNA-seq for mRNA and sRNA, mass spectrometry for protein, and ultra-high pressure liquid chromatography (UPLC) for metabolite and lipid differential regulation in F_0_/F_1_ epididymal white adipose tissue (eWAT), liver, sperm and serum. **(c)** Body weights of F_0_ LF (n=6), F_0_ HF (n=13; n=6 for 15-24 weeks of age), F_0_ HF-LF (n=7, *left*) and F_1_ (F_0_ LF, n=18), F_1_ (F_0_ HF, n=25) and F_1_ (F_0_ HF-LF, n=21, *right*) male C57BL/6N mice. **(d)** Blood glucose during intraperitoneal glucose tolerance test in F_0_ LF (n=6), F_0_ HF (n=6) and F_0_ HF-LF (n=7, *left*) and F_1_ (F_0_ LF, n=10), F_1_ (F_0_ HF, n=10) and F_1_ (F_0_ HF-LF, n=10, *right*) male C57BL/6N mice. **(e)** Blood glucose during intraperitoneal insulin tolerance test in F_0_ LF (n=6), F_0_ HF (n=6) and F_0_ HF-LF (n=7, *left*) and F_1_ (F_0_ LF, n=9), F_1_ (F_0_ HF, n=10) and F_1_ (F_0_ HF-LF, n=10, *right*) male C57BL/6N mice. **(f)** Relative eWAT weights in F_0_ LF (n=6), F_0_ HF (n=6) and F_0_ HF-LF (n=7, *left*) and F_1_ (F_0_ LF, n=10), F_1_ (F_0_ HF, n=10) and F_1_ (F_0_ HF-LF, n=7, *right*) male C57BL/6N mice. **(g)** Serum leptin levels measured by ELISA in F_0_ LF (n=6), F_0_ HF (n=6) and F_0_ HF-LF (n=7, *left*) and F_1_ (F_0_ LF, n=8), F_1_ (F_0_ HF, n=8), and F_1_ (F_0_ HF-LF, n=8, *right*) male C57BL/6N mice. **(h)** Fractional area size distribution of eWAT adipocytes in **(i)** F_0_ LF (n=6), F_0_ HF (n=6) and F_0_ HF-LF (n=6, *left*) and F_1_ (F_0_ LF, n=6), F_1_ (F_0_ HF, n=6) and F_1_ (F_0_ HF-LF, n=6, *right*) male C57BL/6N mice. **(i,j)** Representative haematoxylin and eosin staining of eWAT of (**i**) F_0_ LF, F_0_ HF and F_0_ HF-LF and **(j)** F_1_ (F_0_ LF), F_1_ (F_0_ HF), and F_1_ (F_0_ HF-LF) male C57BL/6N mice. **(k,l)** Visualisation of high-confidence STRING interaction networks of proteins downregulated in (**k**) F_0_ LF (n=6) vs F_0_ HF (n=6) and (**l**) F_1_ (F_0_ LF, n=6), F_1_ (F_0_ HF, n=6) eWAT (*top*) and GOBP enrichment (*bottom*). Two-tailed Student’s t-test **(c-e,h)** for each timepoint and adipocyte sizes or one-way ANOVA followed by Tukey’s multiple comparisons **(f-h)** were used for statistical analysis. Data are presented as mean ± standard error with individual values shown for n≤10. P-Values are indicated in the panel or represented as letters (**a**, *p*<0.05, F_0_ LF versus F_0_ HF or F_1_ (F_0_ LF) versus F_1_ (F_0_ HF); **b**, *p*<0.05, F_0_ HF versus F_0_ HF-LF or F_1_ (F_0_ HF) versus F_1_ (F_0_ HF-LF); **c**, *p*<0.05, F_0_ LF versus F_0_ HF-LF or F_1_ (F_0_ LF) versus F_1_ (F_0_ HF-LF)).

When phenotyping F_0_ and male F_1_ animals for metabolic traits commonly associated with obesity and type 2 diabetes (T2D), we found that HF increased and ensuing HF-LF feeding reduced body weight to levels seen in F0 LF mice, whilst F_1_ progeny were indistinguishable in terms of body weight (**Fig. 1c**). Intriguingly, male obesity impaired glucose tolerance (**Fig. 1d, Fig.S2a**) and insulin sensitivity (**Fig. 1e)** not only in F_0_ HF mice, but also in their F_1_ (F_0_ HF) offsprings without changes in circulating F_1_ insulin (**Fig.S2b**) and no alterations in liver and brown adipose tissue mass (not shown). In contrast, we found that obesity increased iWAT (**Fig.S2c**) and eWAT (**Fig. 1f**) masses and elevated serum levels of leptin, an important adipokine reflecting adipocyte size and numbers, in F_0_ and F_1_ mice (**Fig. 1g**). Adipocyte size determination using haematoxylin/eosine (H/E) staining of formalin-fixated, paraffine-embedded adipose sections demonstrated comparable shifts towards increased eWAT adipocyte sizes in F_0_ and F_1_ (**Fig. 1h-j**), suggesting that paternal obesity might promote lipid accrual/lipolysis as well as metabolic processes governing adipocyte health by similar molecular mechanisms in both generations and via this mechanism contribute to glucose intolerance in F_0_ and male F_1_. When phenotyping female offsprings of lean, obese and weight-regressed F_0_ fathers, we observed no changes in F_1_ weight (**Fig. S2d**), mild glucose intolerance (**Fig.S2e,f**) and only trends towards changes in insulin sensitivity (**Fig.S2g**). In support of overall mild impairments in female offsprings, random fed glycemia (**Fig.S2h**) and organ weights (**Fig.S2i**) were also unchanged in female progeny. *Thus, male obesity and weight loss reversibly affect glucose tolerance in F*_1_*, with metabolic effects predominantly observed in male progeny*.

For comprehensive insights into molecular processes affected by paternal obesity and weight loss, and those governing intergenerational effects, we performed bulk tissue mass spectrometry to determine differentially abundant proteins in F_0_ and F_1_ eWAT. We observed 980/431 proteins increased in F_0_ HF compared to F_0_ LF mice and in F_1_ (F_0_ HF) compared to F_1_ (F_0_ LF) mice. Conversely, 632/305 proteins were less abundant comparing F_0_ HF to F_0_ LF mice (**Fig.S3a**) and F_1_ (F_0_ HF) compared to F_1_ (F_0_ LF) mice (**Fig.S3b**). Using the STRING database (https://string-db.org/) followed by Gene Ontology (GO) analysis of overrepresented biological processes (GOBP), we intriguingly saw that F_0_ HF mice (**Fig. 1k**) and F_1_ (F_0_ HF) progeny (**Fig. 1l**) shared the same downregulated, mitochondria-associated GOBPs such as Oxphos, adenosine triphosphate (ATP) synthesis, and regulation of ETC subunits. Examples of proteins similarly repressed in F_0_ HF and F_1_ (F_0_ HF) eWAT included components of ETC Complex I like NADH dehydrogenase 1 alpha subcomplex subunit 8 (NDUFA8) and NADH dehydrogenase iron-sulfur protein 6 (NDUFS6), of Complex IV like Cytochrome C Oxidase Subunit 6B1 (COX6B1), of mitochondrial membrane-intrinsic lipid carriers such as *Carnitine Palmitoyltransferase 1A* (CPT1A) and of mitochondrial protein translocation such as *Translocase Of Outer Mitochondrial Membrane 20* (TOMM20). In contrast, upregulated GOBP terms in F_0_ HF mice (**Fig.S3a**) and their F_1_ (F_0_ HF) progeny (**Fig.S3b**) included unrelated GOBP terms like protein translation, Golgi-mediated transport and secretion as well as protein localisation. Importantly, abundance of metabolite– (**Fig.S4a-c**) and lipid-based (**Fig.S4d,e**) biomarkers that can reflect systemic mitochondrial dysfunction, for instance tricarbon cycle acid (TCA) intermediates like citrate, malate, fumarate, alpha-ketoglutarate and citrate, were reduced by paternal obesity and restored by weight loss (**Fig.S4f**) and mitochondrial lipid carriers like (acetylated) carnitine (**Fig.S4g**) were reduced. Furthermore, saturated, short-chain C:16:0/C:18:0 ceramides, i.e., lipid species reported to disrupt mitochondrial function and elicit mitochondrial ER stress in obese adipocytes^53,54^ were induced, whilst mito-protective long-chain C18:1/C24:1 ceramides were reduced in F_0_ HF mice (**Fig.S4h**). *Thus, male obesity downregulates proteins associated with mitochondrial processes in obese F*_0_ *and F*_1_ *offsprings that correlate with reduced levels of mitochondrial function and lipid catabolism, likely reflecting a system-wide hereditary form of mitochondrial dysfunction, also in adipose tissue*.

### Paternal obesity negatively affects expression of mitochondria-associated miRNA-mRNA networks

We next turned our attention to in detail study gene-regulatory processes at the messenger and miRNA level that might underly the observed repression of mitochondria-associated proteins in F_0_ and F_1_ adipose tissue, and to this end performed mRNA-seq of eWAT and liver of F_0_ and F_1_ groups and performed miRNA and mRNA differential expression analysis, specifically focussing on mitochondrial processes. Comparing F_0_ HF to F_0_ LF eWAT using adjusted p-Values (pV_adj_) ≤0.05 we detected yielded 3,265 up-regulated and 2,890 down-regulated genes (**Fig. 2a**). When comparing F_1_ (F_0_ HF) to F_1_ (F_0_ LF) mice we detected no significantly regulated genes when accounting for multiple testing (pV_adj_). Yet when specifically interrogating differential expression to reflect the more variable transcriptional effects in F_1_ mice, we found 766 up-regulated and 893 down-regulated genes using unadjusted p-Values (pV) ≤ 0.05 (**Fig. 2b**). When performing gene set enrichment analysis (GSEA) of affected GOBP we found upregulated processes linked to cell proliferation and immune activation that likely reflect adipocyte hyperplasia and inflammation, thus cellular processes commonly associated with obesity, and GOBP associated with germ cell and cilial function in F_1_. Importantly, we confirmed our proteomic findings and detected overrepresented GOBP related to lipid catabolism, ATP synthesis, OxPhos and ETC downregulated in adipose tissue of F_0_ HF mice (**Fig. 2c**) and their F_1_ offsprings (**Fig. 2d**) and, when interrogating expression of mRNAs encoding mitochondrial ETC I-V, we confirmed the repression of nuclear-expressed mitochondrial constituents at the mRNA level (**Fig. 2a,b**). Importantly, aberrant mitochondrial effects were to large extends reversible by weight loss, as comparing F_0_ HF to F_0_ HF-LF eWAT yielded 2,687 up-regulated and 2,237 down-regulated using pV_adj_ (**Fig.S5a**), whilst comparing F_1_ (F_0_ HF) to F_1_ (F_0_ HF-LF) eWAT yielded 547 up-regulated and 732 down-regulated genes without adjusting for multiple testing, (**Fig.S5b**) and GOBP restored by weight loss also fell into genes linked to mitochondrial function (**Fig.S5c,d**).

**Figure 2:**
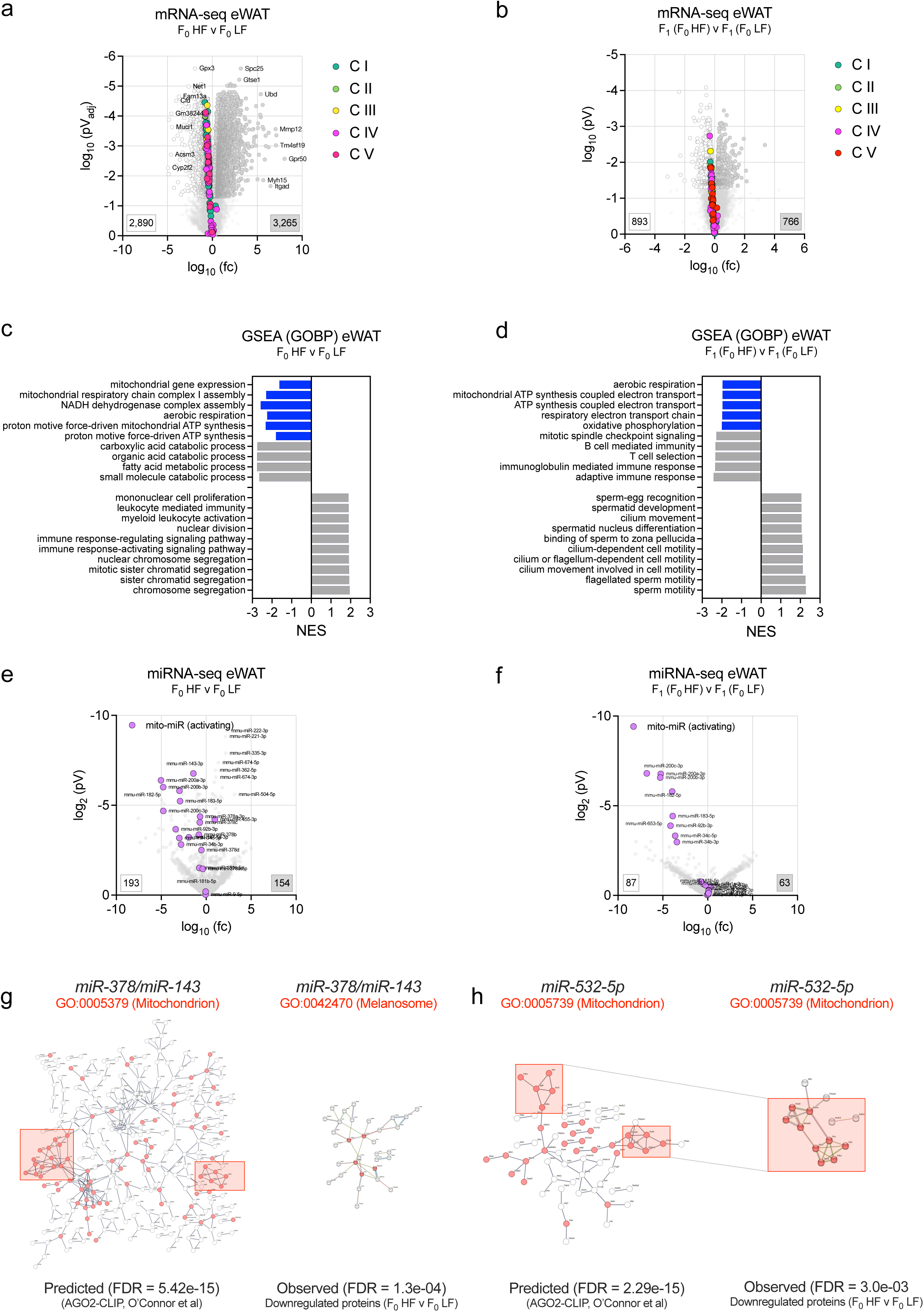
Paternal obesity dysregulates miRNA:mRNA gene networks spurring intergenerational mitochondrial dysfunction. (**a,b**) Volcano plot of significantly up– (grey) and down-regulated (white) mRNAs in eWAT from **(a)** F_0_ HF versus F_0_ LF with 3,495 upregulated and 1,639 downregulated mRNAs and (**b**) F_1_ (F_0_ HF) versus F_1_ (F_0_ LF) with 1,284 up– and 702 down-regulated mRNAs. Plots show log10 transformed fold-changes (fc) and log10-transformed **(a)** adjusted or **(b)** non-adjusted p-Values of mRNA changes. Mitochondrial complex I-V genes are annotated in the plot. **(c,d)** Gene Set Enrichment Analysis and gene ontology enrichment of biological processes (GOBP) of differentially expressed mRNAs from **(a,b).** GOBP linked to mitochondrial respiration, ATP synthesis and Complex I assembly are marked with blue bars. **(e,f)** Volcano plot of significantly regulated miRNAs in eWAT from **(e)** F_0_ HF versus F_0_ LF with 154 up– and 193 down-regulated miRNAs and (**f**) F_1_ (F_0_ HF) versus F_1_ (F_0_ LF) with 63 up– and 87 downregulated miRNAs. Plots depict log10 transformed fc and log10-transformed p-Values of expression changes. Mitochondria-activating miRNAs are annotated in pink. **(g,h)** STRING interaction networks of **(g)** *miR-378/miR-143* and **(h)** *miR-532-5p* targets predicted by AGO2-CLIP (*left*) and network subset for eWAT proteins downregulated when comparing F_0_ LF (n=6) versus F_0_ HF (n=6) male C57BL/6N mice. Submodules containing mitochondrial proteins are marked by a red square and proteins marked by red dots are AGO2-CLIP predicted *let-7* targets and downregulated in F0 HF versus F_0_ LF eWAT.

Given that miRNAs are important (rheostatic) components in post-transcriptional gene network and have previously been linked to mitochondrial maturation and function^55^, albeit not in adipocytes, we next wondered whether concerted down-regulation of mitochondrial programs in obese eWAT might (partially) be instigated by altered expression of mito-miRs^1^’ like *let-7*^2^, *miR-378*^3^, *miR-181*^4^, *miR-143*^5^ that functionally support mitochondrial formation and oxidative metabolism. Thus, when conducting small RNA (sRNA)-seq to determine differential miRNA expression, we found that F_0_ obesity repressed mito-miRs that positively affect mitochondrial biogenesis and function, for instance *miR-143-3p*^56^, *miR-181a-2-3p*^57,58^, *miR-182*^59^, *miR-200*^60^ and *miR-378a-3p*^61^ in eWAT of F_0_ HF (**Fig. 2e**) and F_1_ (F_0_ HF, **Fig. 2f**) mice, whereas mitochondria-inactivating *miR-532*^62^ was induced in F_0_ HF (**Fig. 6c**) and F_1_ (F_0_ HF, **Fig. 6d**) eWAT, suggesting that miRNA dysregulation contributed to mitochondrial impediments observed in obese F_0_ eWAT. Integration of miRNA-mRNA NGS datasets with adipocyte-specific miRNA-mRNA-Argonaute 2 interactomes using AGO-HITS-CLIP^6^ revealed that *miR-378 and miR-143*, despite being predicted, did not coinidce with downregulation of their mitochondrial protein targets (**Fig. 2g**), whilst *miR-532-5p* was both predicted to target and observed to coincide with mitochondrial proteins repressed in F0 obesity, arguing for roles of *miR-532,* but not *miR-143/378* in driving mitochondria dysfunction in F_0_ HF eWAT (**Fig. 2h**). In line with body weight loss-driven mRNA transcriptional responses (**Fig.S6a,b**), miRNA alterations were highly dynamic, and weight loss restored expression of mitochondria-promoting in F_0_ HF (**Fig.S5e**) and F_1_ (F_0_ HF, **Fig.S5f**) mice. *Thus, F*_0_ *obesity causes reversible repression of mitochondrial and lipid catabolic genes, and miRNA-mRNA network analysis implicates obesity-regulated mito-miRs in this process*.

### The specificity of obesity-induced heritable epigenetic mitochondrial alterations to adipose tissue

We were in the following interested to address how confined the observed heritability of mitochondrial dysfunction was i. as a cellular and metabolic process and to ii. adipose tissue and performed mRNA-seq to assess the heredity of other obesity-associated processes such as chronic, low-grade inflammation (meta-inflammation). Focussing on expression of gene markers of classically (M1) and alternatively activated (M2) monocytes/macrophages, cell types that are linked to meta-inflammation-driven adipose tissue dysfunction^33^, we found that, as expected, mRNAs encoding M1 markers such as *Interleukin 1b (IL1b)*, *Monocyte chemoattractant protein-1 (Mcp1/Ccl2)* and *Tumor Necrosis Factor Alpha (TNF)*; but also M2 markers like *Arginase 1 (Arg1)*, *Mannose Receptor C-Type 1 (Mrc1)*, *C-Type Lectin Domain Containing 10A (Clec10a)* and *Resistin Like Beta (Retnla)* were induced in F_0_ HF, but not F_1_ eWAT (**Fig.S6e,f**), emphasising that a meta-inflammatory tone is not passed on from F_0_ HF mice.

Given the dysfunction in a specialised cellular process like mitochondrial respiration in eWAT, we next aimed to juxtapose adipose tissue with transcriptional effects in liver, an organ essential for glucose and lipid homeostasis: Analysis of mRNAseq data using multidimensional scaling (MDS) plots showed that F_0_ LF segregated from F_0_ HF and F_0_ HF-LF, arguing for mostly irreversible effects of obesity in F_0_ liver (**Fig.S7a**) and modest effects of F_0_ obesity on F_1_ liver transcriptomes (**Fig.S7b**). When comparing F_0_ HF to F_0_ LF livers, we found 2,474 up– and 2,612 down-regulated genes with pV_adj_ ≤0.05 (**Fig.S7c**), whereas comparing F_1_ (F_0_ HF) to F_1_ (F_0_ LF) without adjusting for multiple testing (pV ≤0.05) yielded 716up– and 546 down-regulated genes (**Fig.S7d**). Gene Set Enrichment analysis (GSEA)^63^ of differentially regulated mRNAs identified lipid metabolism and organelle formation as induced, whereas repressed terms included lipoprotein trafficking, endoplasmic reticulum and ribosome, yet no transcriptional evidence for altered mitochondrial processes in liver of obese mice and their F_1_ progeny (**Fig.S7e,f**), emphasising the specificity of mitochondrial alterations in obese fat.

The non-hereditary effects of paternal obesity were supported by histological evidence, and pathologist assessment of H/E (morphology), Sirius Red (collagen/fibrosis) and anti-CD45 (leukocytes) staining revealed important hallmarks of metabolic dysfunction-associated steatosis liver disease (MASLD) like micro-/macro-vesicular steatosis, fibrosis and locular inflammation in F_0_ HF mice, but not lean F_0_ or F_1_ descendants (**Fig.S8a-h**). Targeted analysis of expression changes in genes linked to hepatic lipid metabolism and meta-inflammation such as *Adhesion G Protein-Coupled Receptor E1 (Adgre, F4/80)*, *Apolipoprotein A4 (Apoa4), Apolipoprotein E (Apoe), Low Density Lipoprotein Receptor (Ldlr), Peroxisome Proliferator Activated Receptor Gamma (Pparg), Cluster of Differentiation 36 (Cd36)* and *Sterol Regulatory Element Binding Transcription Factor 1 (Srebf/Srebp1)* confirmed that induction was confined to F_0_ HF livers (**Fig.S8i**). Furthermore, mitochondria-associated genes that were activated in obesity, presumably counteracting lipid accumulation in MASLD, for instance *Carnithine Palmitoyltransferase 1A (Cpt1a)*, *Carnithine O-Acetyltransferase (Crat), Acyl-Coa-Oxidase (Acox1)* and *Acetyl-CoA-Acetyltransferase (Acat1)* were induced in livers from F_0_ HF, but not F_1_, mice. *Thus, combinatorial analyses of transcriptional and histological changes argue against prominent effects of paternal obesity onto F*_1_ *gene expression and mitochondrial processes in hepatic cell types*.

### Paternal obesity causes qualitatively similar alterations in eWAT and sperm of obese male mice

Spermic sncRNA were reported to be subjected to dietary cues such as high-fat and –sugar diets in mice and humans^22^; and it is therefore conceivable that HF feeding and weight loss alter spermic sncRNA levels that can then, following oocyte fertilisation, be passed on to fertilised oocytes (zygotes) as shown recently ^23^. By changing sRNA composition in zygotes, sncRNAs thus have the unique ability to post-transcriptionally affect decision points in embryogenesis, thereby contributing to the observed glucose intolerance and mitochondrial protein repression observed in F_1_ offsprings sired by F_0_ HF bucks. A particular aspect of sperm transcriptomes is the complex profile of sncRNAs involving miRNAs, ribosomal RNAs (rRNAs), PIWI-interacting RNAs (piRNAs), (mitochondrial) transfer RNAs ((mt)-tRNAs) and (mt)-tRNA fragments ^26^ that, as described by Tomar et al.^23^, are affected by mitochondrial dysfunction in obesity and, particularly relevant in this setting, are physically exchanged between sperm and oocytes during fertilisation, a scenario also conceivable for other types of mitochondria-associated sncRNAs, for example mito-miRs. To the map the effects of paternal obesity and weight loss on spermic sncRNAs and to gain insights into sRNA-driven metabolic adaptations in F_0_ HF sperm, we next isolated motile spermatozoa using swim-up assays and charted differential spermic sncRNAs responses as consequence of body weight gain and losses and found sperm harboured complex sncRNA pools of thousands of (mt)-tRNAs, piRNAs, rRNAs and miRNAs (**Fig. 3a-c**). In addition to effects elicited by obesity and/or LF feeding at the single sncRNA level, we observed that (mt)-tRNAs and miRNAs as entire RNA biotype were repressed by HF feeding. As a prior study from Tomar et. al^23^ had shown effects on mt-RNA in sperm after HF feeding, this was investigated further: Mt-tsRNA fragments were found for nine mt-tRNAs (*mt-Tl1*, *mt-Tq*, *mt-Tm*, *mt-Ts1*, *mt-Tr*, *mt-Th*, *mt-Ts2*, *mt-Te*, *mt-Tp*), where mean cpm above 10 was observed in *mt-Tm*, *mt-Ts1*, *mt-Ts2* and *mt-Th* (**Fig. 9a**), but only 5’ fragments of *mt-Ts-1* were significantly changed in sperm from F_0_ HF and F_0_ HF-LF diet (**Fig. 9b**). As miRNAs are abundant in somatic cell types and sperm, harbour the possibility of horizontal transfer between them, and exert gene-regulatory functions within both cellular compartments, we turned our attention to miRNA regulation in sperm. We found that F_0_ obesity and weight loss caused a coordinated and inverse (i.e., downregulation and upregulation, respectively) effects on miRNome regulation (**Fig. 3d,e**), a finding previously observed by our labs ^51^ and others^49,50,64^ for obese adipose tissues; in the latter case at least partially due reduced abundance of DICER1. Intrigued by the repression of miRNAs in two functionally distinct, yet anatomically adjacent tissues and cell types like eWAT and sperm, we asked if differential miRNA responses were qualitative similar in both organ which, if present, would support the possibility of physical miRNA exchange between eWAT adipocytes and sperm as shown for other testicular cell types and sperm^21^: Indeed, upon comparison of lean, obese and weight-regressed sperm and eWAT miRNAs, we found highly congruent responses across both compartments with activating mito-miRNAs like *miR-378* and *miR-143* being down-regulated, whilst inactivating mito-miRs of the *let-7* family up-regulated in obese F_0_ eWAT and sperm (**Fig. 3f)**. Importantly, when broadly mapping the regulation of *let-7* family isoforms, we found mild, but synchronous inductions of *let-7* in F_0_ eWAT and sperm and F_1_ eWAT (**Fig. 3g-3i**). As increases of *let-7c* were reported in intergenerational rat studies by us ^8^ and others^65^ and because *let-7* impairs embryo implantation and glucose homeostasis when overexpressed in mice ^66,67^, we focussed on *let-7* isoforms for functional follow-up studies in germ cells and adipocytes. *Thus, paternal obesity triggers similar alterations in adipose tissue and sperm miRNAs and let-7 isoforms are consistently induced across cell types and generations*.

**Figure 3:**
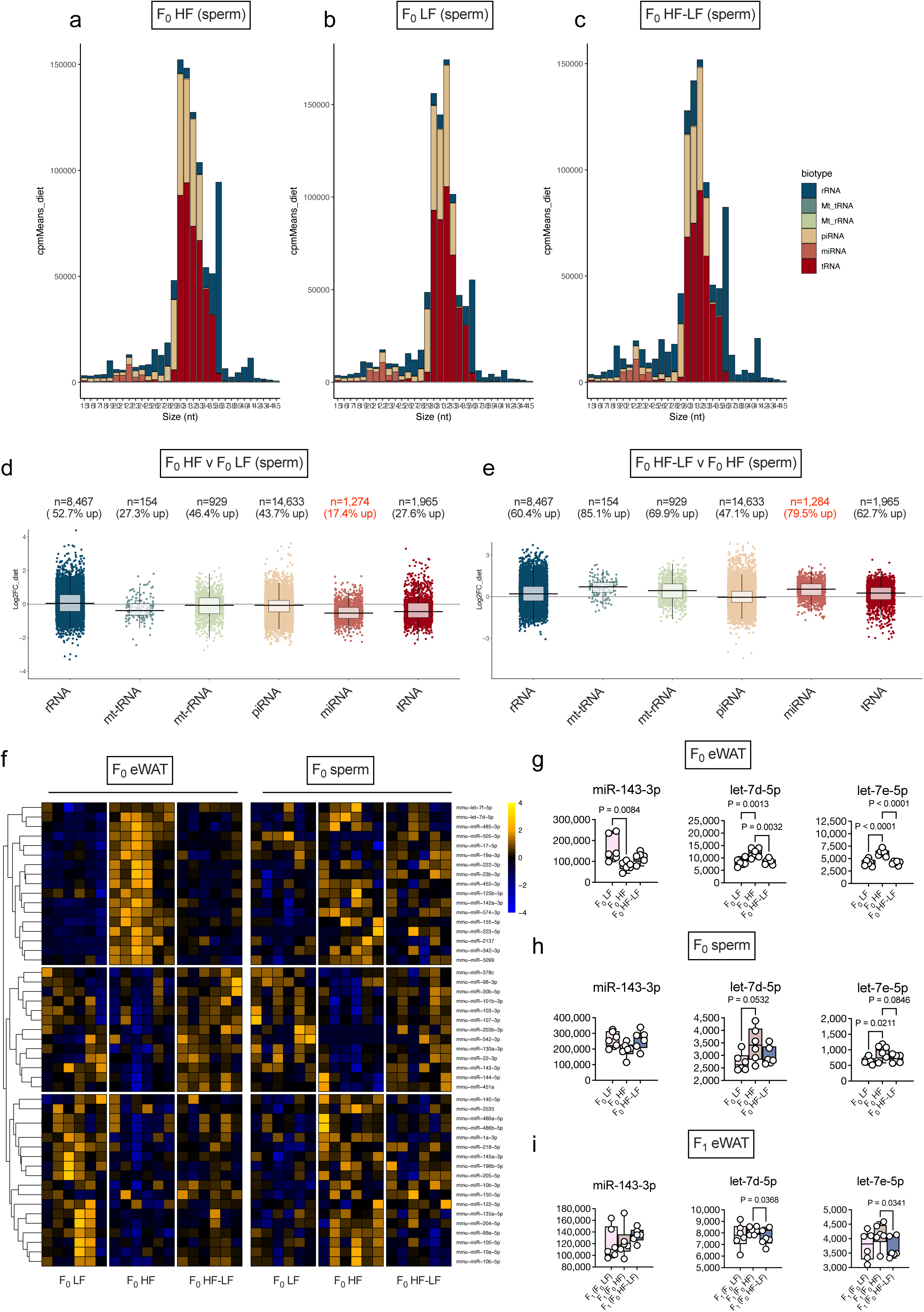
Obesity reversibly affects sperm sncRNomes and induces *let-7* in eWAT and sperm. **(a)** Size distribution on nucleotide length of biotypes present in F_0_ HF-LF sperm. Data presented is mean cpm per diet. **(b)** Size distribution on nucleotide length of biotypes present in F_0_ HF sperm. Data presented is mean cpm per diet. **(c)** Size distribution on nucleotide length of biotypes present in F_0_ LF sperm. Data presented is mean cpm per diet. **(d)** Fold change of F_0_ HF vs F_0_ LF sncRNA in sperm. Each points represents one sequence. **(e)** Fold change of F_0_ HF-LF vs F_0_ HF sncRNA in sperm. Each points represents one sequence. Dark blue= Mitochondrial tRNA, dark green=piRNA, green=ribosomal RNA, light green =long non-coding RNA, yellow=mitochondrial ribosomal RNA, orange= miRNA, dark orange=protein coding, red= miscellaneous RNA, dark red= tRNA, grey=unannotated. **(f)** Heatmap depicting scaled abundances of miRNAs in F_0_ LF, F_0_ HF and F_0_ HF-LF (n=6/group) eWAT and sperm as determined by sRNA-seq. **(g-i)** Abundance of *miR-143-3p*, *let-7d-5p* and in F_0_ LF (n=6), F_0_ HF (n=6) and F_0_ HF-LF (n=6) eWAT and sperm. One-way ANOVA followed by Tukey’s multiple comparisons (g-i) were used for statistical analysis. Data are presented as mean ± standard error with individual values shown for n≤10. P-Values are indicated in the panel.

### Zygotic delivery of let-7 causes glucose intolerance and mitochondrial impairment in sired mice

Prompted by profound and reversible effects of male obesity and weight loss on sperm miRNomes and *let-7* in particular, as environmentally and artificially induced (e.g., elicited by miRNA microinjection) alteration in zygote miRNA (pools) trigger abnormalities in progeny^25,68^, and because embryonic trajectories require maturation of pre-into mature miRNAs^69^, we tested the functional relevance of *let-7* gain-of-function in one-cell embryos for F_1_ phenotypes. We focussed on *let-7d-5p* and *let-7e-5p* given that these exhibit the most concordant increases amongst *let-7* family members in F_0_ and F_1_ eWAT and sperm (**Fig. 3g-i**) and because *let-7* has before been connected to embryonic cell fate decisions^70,71^. To test if zygotic delivery of *let-7* corresponding at levels seen in few obese spermatozoa triggers phenotypical and metabolic changes in mice sired from *let-7* microinjected embryos (F_MI_ generation) and whether these would resemble mice sired by F_0_ HF fathers, we injected physiological amounts (50 fmol, i.e., *let-7* found in 5-10 spermatozoa) of *cel-mir-67* mimic as non-mammalian (*C.elegans*) control, and mimics for *let-7d-5p* and *let-7e-5p* by microinjection into zygotes conceived from lean males and females C57BL/6N mice using standard protocols^29^ (**Fig. 4a**). We injected 126/138/106 zygotes with *cel-mir-67*, *let-7d-5p* and *let-7e-5p*, but observed no discernible difference in implantation success, litter sizes and number of pregnancies in F_MI_ mice (**Fig.S10**), arguing against roles of *let-7* for zygote viability and implantation. Next, we analysed at least 19 pups per group from 3-4 pregnancies but observed no overt differences in postnatal body weight trajectories (**Fig. 4b**). Despite similar weight, we detected impaired glucose tolerance (**Fig. 4c,d**), insulin resistance (**Fig. 4e**) and elevated random-fed glycemia (**Fig. 4f)** in mice sired by *let-7* injected F_MI_ embryos, whilst eWAT and liver weights were indistinguishable between F_MI_ groups (**Fig. 4g**). Importantly, when performing mRNA-seq in eWAT of F_MI_ mice, despite mild changes, we found that *let-7* down-regulated mitochondrial processes in F_MI_ mice (**Fig. 4h**). *Thus, zygotic delivery of let-7 at levels corresponding to few F*_0_ *HF sperm cells, causes glucose intolerance and aberrant mitochondrial gene regulation in mice derived from these embryos*.

**Figure 4:**
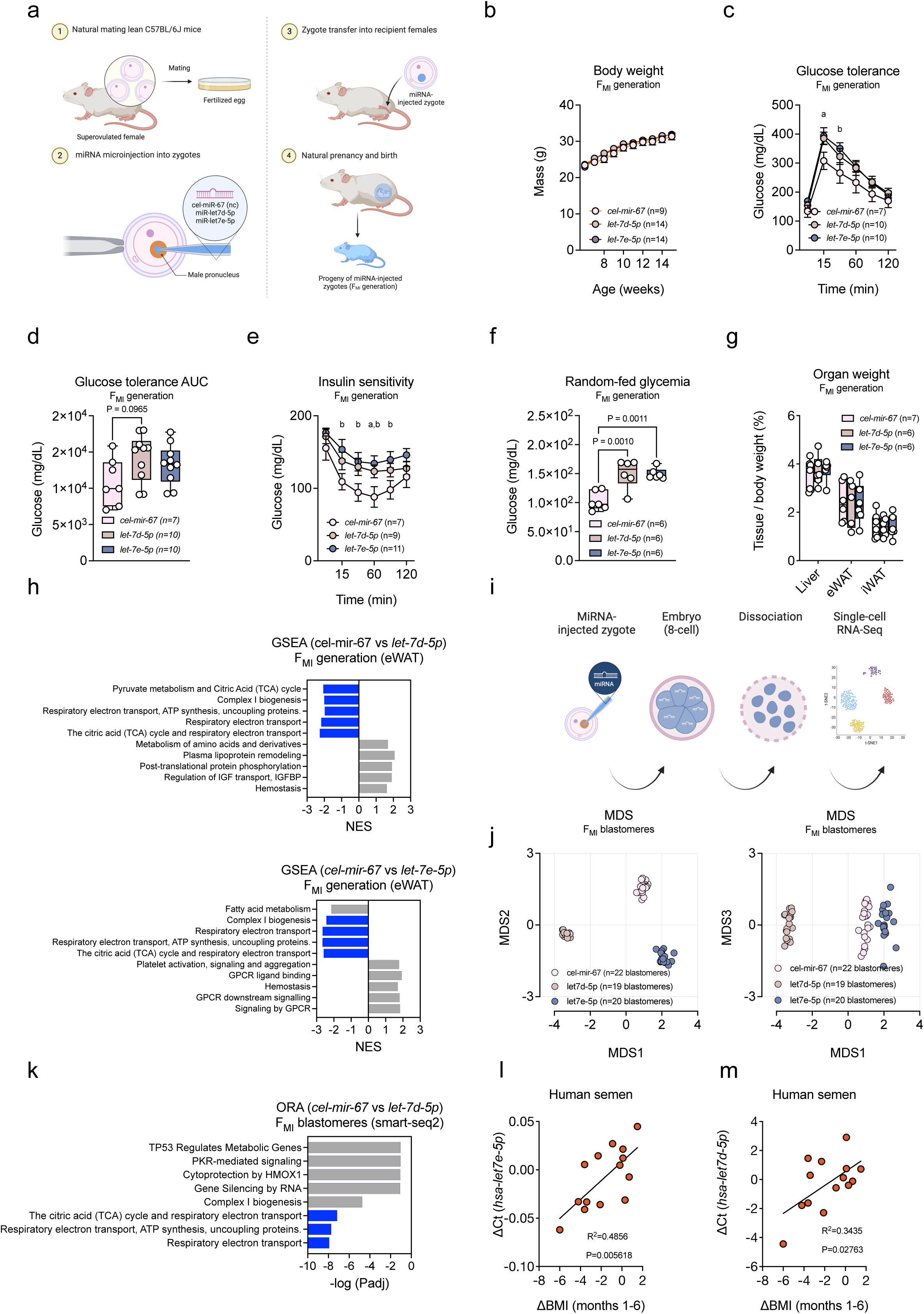
Zygotic delivery of *let-7* causes glucose intolerance in sired F_MI_ mice and impairs embryonic mitochondria. (**a**) Illustration of experimental workflow for zygotic miRNA injection and phenotypic characterisation of sired offsprings (F_MI_ mice). **(b)** Body weights of male F_MI_ mice sired from with *cel-mir-67* (n=9), *let-7d-5p* (n=14) and *let-7e-5p* (n=14) injected zygotes. **(c,d)** Blood glucose levels **(c)** and glucose AUC **(d)** during intraperitoneal glucose tolerance test in male F_MI_ mice sired from *cel-mir-67cel-mir-67* (n=7), *let-7d-5p* (n=10) and *let-7e-5p* (n=10) injected zygotes. **(e)** Blood glucose levels during intraperitoneal insulin tolerance test in male F_MI_ mice sired from *cel-mir-67* (n=7), *let-7d-5p* (n=9) and *let-7e-5p* (n=11) injected zygotes. **(f)** Random-fed blood glucose at indicated in male F_MI_ mice sired from *cel-mir-67* (n=6), *let-7d-5p* (n=6) and *let-7e-5p* (n=6) injected zygotes **(g)** Relative weights of indicated organs from male F_MI_ mice sired from *cel-mir-67* (n=7), *let-7d-5p* (n=6) and *let-7e-5p* (n=6) injected zygotes. **(h)** GSEA-GOBP analysis of eWAT of male F_MI_ mice sired from *cel-mir-67* (n=5), *let-7d-5p* (n=6) and *let-7e-5p* (n=6) treated zygotes. GOBP linked to mitochondrial respiration, ATP synthesis and Complex I assembly are marked with blue bars. **(i)** Illustration of workflow to obtain single 8-cell stage blastomeres derived from zygotes injected with *cel-mir-67*, *let-7d-5p* and *let-7e-5p* subjected to smart-seq2 single-cell sequencing. **(j)** Multidimensional scaling plots of single-cell RNAseq profiles in 8-cell stage blastomeres obtained from *cel-mir-67* (n=22 blastomeres derived from n=3 zygotes), *let-7d-5p* (n=19 cells blastomeres from n=3 zygotes) and *let-7e-5p* (n=20 blastomeres derived from n=4 zygotes). **(l,m)** Correlation plot and regression analysis between differential BMI loss (ΔBMI) and expression changes (ΔCt) in human semen (**l**) *LET-7D-5P* and (**m**)*LET-7E-5P.* Two-tailed Student’s t-test **(c,e)** per timepoint, one-way ANOVA followed by Tukey’s multiple comparisons **(d,g)** or Pearson’s regression analysis **(l,m)** were used for statistical analysis. Data are presented as mean ± standard error with individual values shown for n≤10. P-Values are indicated in the panel or represented as letters (**a**, *p*<0.05, *cel-mir-67* versus *let-7d-5p*, **b**, *p*<0.05, *cel-mir-67* versus *let-7e-5p*).

### Obesity-associated let-7 rewires mitochondrial metabolism during embryogenesis

As transfer of miRNA perturbation in sperm to oocytes can have deleterious consequences and give rise to incompletely developed embryos ^20,72,73^, and as zygotes and blastocysts have the capacity to differentiate into virtually all cell types, (early) embryogenesis represents an important window of developmental plasticity that might affect F_MI_ phenotypes if perturbed by miRNA delivery, despite the physiological levels of miRNA delivery and absence of embryonic lethality. Nonetheless, it is conceivable that alterations in zygotic miRNAs still entails far-reaching consequences in sired mice due to a combination of altered developmental, epigenetic and metabolic processes^73^. Based on the specific mitochondrial alterations, but absence of increased *let-7* levels in F_MI_ eWAT, we next hypothesized that *let-7d-5p* and *let-7e-5p* could carry out important roles in cell fate decisions during embryogenesis which will indirectly contribute to the glucose intolerance in F_MI_ pups via altered embryonic development. To test this experimentally, we repeated zygotic *let-7* injections but allowed miRNA-injected embryos to divide thrice until the 8-cell blastomere stage. We dissociated embryos into single blastomeres and performed single-cell RNA-Seq (scRNA-seq) using Smart-seq2^74^ (**Fig. 4i**). Remarkably, we found that after three cell divisions, miRNA-injected blastomeres were still transcriptionally distinct, reflecting the profound transcriptional and developmental consequences of inducing *let-7* in zygotes (**Fig. 4j**). When performing GSEA of *let-7*-dependent gene sets, we corroborated that *let-7d-5p* repressed terms linked to mitochondrial processes like ATP synthesis, TCA cycle and ETC (**Fig. 4k**). Intergenerational programming in isogenic rodents has been described before, yet it is elusive if these effects are translatable to species with highly different reproductive apparatuses, genetic backgrounds, and timescales of obesity development like humans. Although epidemiological evidence^4,75^ suggests heritable effects of overfeeding and malnutrition in humans (e.g., the famous ‘Dutch Hunger Winter’^76^), causal evidence for epigenetic inheritance is hard to separate from cultural and ecological inheritance in humans^2^. As it is becoming appreciated that nutritional and metabolic cues shape human sncRNA expression^22^, we tested effects of weight change on semen *let-7* in a small cohort of n=15 obese men (BMI, 39.49 ± 1.80) undergoing voluntary weight loss^77^ and quantified *hsa-let-7* using TaqMan qPCR. Intriguingly, we found that *hsa-let-7d-5p* and *hsa-let-73-5p* followed the trends observed for weight regulation, suggesting similar correlations between metabolic improvement and *let-7* reductions in human semen (**Fig. 4l,m**). *Thus, obesity-associated let-7d affects murine embryogenesis by impairing mitochondrial gene regulation and similar processes might be occurring following weight loss in obese humans*.

### Adipocyte let-7 impairs mitochondrial function via post-transcriptional silencing of DICER1

Beyond F_0_ HF sperm (**Fig. 3g-i**), we found *let-7d-5p* and *let-7e-5p* upregulated in F_0_/F_1_ eWAT, and AGO2-CLIP interactome analyses confirmed that *let-7* gene targets are enriched for genes encoding mitochondrial proteins, that are downregulated by obesity in F_0_/F_1_ eWAT (**Fig. 5a**). For this, and to add to their previously ascribed role in fertilized oocytes, we aimed to dissect the transcriptional and functional consequences of elevated *let-7* in adipocytes, with a specific focus on mitochondrial gene regulation as highly sequence-related *let-7i* can impair commitment of mitochondria-rich adipocytes ^78^. Yet, if roles for *let-7* in mitochondrial formation and function exist in mature adipocytes is unknown. To test this, we delivered by reverse lipofection mimetics for *let-7d-5p* and *let-7e-5p* and *cel-mir-67* into stromal vascular fraction-derived mature epididymal white adipocytes (1°eWA) to mimic obesity-associated increased in *let-7* in eWAT. When interrogating the effects of *let-7* overexpression, we observed that *let-7d-5p* elicited more pronounced effects (1,768 up-regulated, 1,387 down-regulated genes, **Fig. 5b**) than *let-7e-5p* (557 up-regulated, 893 down-regulated genes, **Fig. 5c**) compared to *cel-mir-67*, yet both *let-7* isoforms downregulated processes related to lipid metabolism, Insulin-AKT (Protein Kinase B) signalling and carbohydrate metabolism (**Fig.S11**). Although not overrepresented at the GOBP level, we found rate-limiting genes in glycolysis, TCA, and mitochondrial lipid import like Carnithine Palmitoyltransferase 2 (*Cpt2*), Carnithine O-Acetyltransferase (*Crat*), Hexokinase 2 (*Hk2*), Isocitrate Dehydrogenase 2 (*Idh2*), Pyruvate Carboxylase (*Pcx*), Pyruvate Dehydrogenase Kinase 2 (*Pdk2*) and mitochondrial ATP-magnesium/phosphate antiporter Solute Carrier Family 25 Member 24 (*Slc25a24*, **Fig. 5b,c**; in blue) repressed upon *let-7* delivery. Importantly, we also detected pivotal genes in miRNA maturation such as Ribonuclease III DICER1 *(Dicer1)* and Argonaute RISC Catalytic Complex 2 *(Ago2,* **Fig. 5b,c**; in red). Next, we experimentally validated these predicted *let-7* targets by dual luciferase reporter assays (DLRA)-based miRNA-mRNA 3’UTR interaction studies in HEK293 cells for genes linked to mitochondrial lipid and ATP uptake (*Slc25a24, Cpt2*), glycolysis (*Hk2*, *Pdk2*), TCA activity (*Sdha*, **Fig. 5d**) and miRNA processing (*Dicer1, Ago2*).

**Figure 5:**
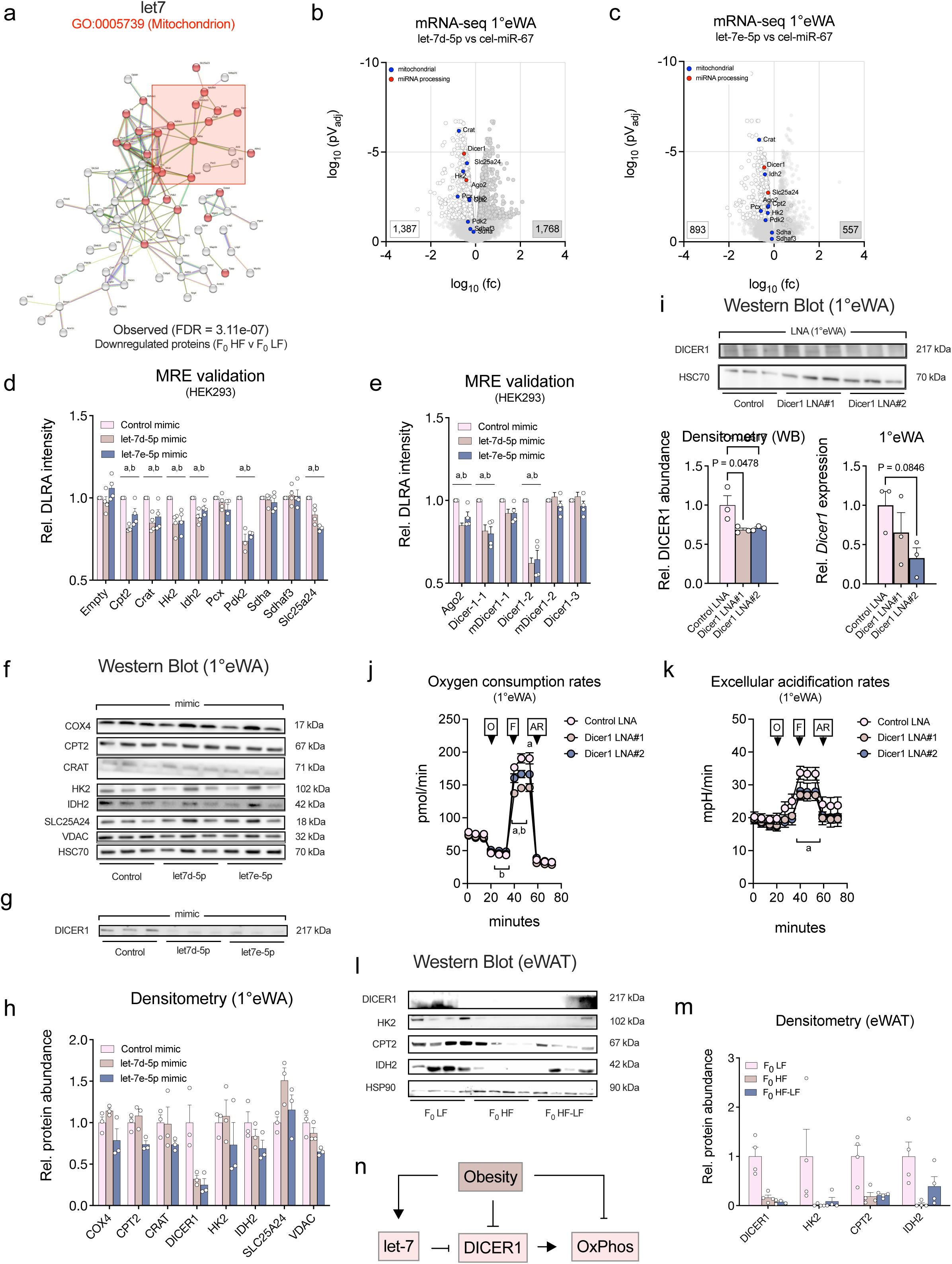
Adipocyte *let-7* blunts mitochondrial function via a novel let-7-DICER1 axis. **(a)** Visualisation of high-confidence STRING interaction networks of *let-7* targets predicted by AGO2-CLIP miRNA:mRNA interactions and subsetted for proteins downregulated comparing F_0_ LF (n=6) versus F_0_ HF (n=6) in eWAT from male C57BL/6N mice. Submodules containing mitochondrial proteins are marked by a red square and proteins marked by red dots are AGO2-CLIP predicted *let-7* targets and downregulated in F0 HF versus F_0_ LF eWAT. **(b,c**) Volcano plot of significantly up-(grey) and down-regulated (white) genes in 1°eWA transfected with (**b**) *let-7-d-5p* versus *cel-mir-67*, yielding 974 up– and 1,295 down-regulated genes and (**c**) *let-7-e-5p* versus *cel-mir-67*, yielding 754 up– and 633 down-regulated genes. Genes connected to mitochondrial regulation are marked by blue, genes involved in miRNA maturation by red dots. **(d)** Relative luciferase activity in HEK293 cells after transfection of pmirGLO dual luciferase reporter assay (DLRA) constructs harbouring wild-type 3′-UTR or exon containing *let-7* miRNA responsive elements (MRE) of indicated mitochondrial genes and (**e**) wild-type and mutated *let-7* MREs in *Dicer1* transfected with 100 nM of *cel-mir-67*, *let-7d-5p* and *let-7d-5p* mimics. Data represent n≥3 independent biological replicates, each performed in n=3 technical replicates. **(f-h)** Western blot analysis of 1°eWA transfected with *cel-mir-67*, *let-7d-5p* and *let-7d-5p* mimics performed using the indicated antibodies detecting **(f)** mitochondrial and **(g)** DICER1 proteins. HSC70 served as loading control. **(h)** Densitometric quantification of (**f,g)** representing n=3 biological replicates. **(i)** Western blot (*top*) and quantitative RT-PCR (*bottom*) quantification of DICER1 in 1°eWA transfected using two independent locked nucleic acid (LNA) inhibitors targeting *Dicer1* mRNA. Data represent n=3 biological replicates, each performed in n=3 technical replicates. **(j,k)** Oxygen consumption rates (**j,** OCR) and extracellular acidification rates (**k**, ECAR) in 1°eWA transfected with 100 nM of *negative control LNA*, *Dicer1* LNA1 and *Dicer1* LNA2 and stimulated with oligomycin (O), FCCP (F) and Antimycin A plus Rotenone (A/R). Experiments are representative of n≥3 independent biological replicates. **(l)** Western blot analysis of eWAT from F_0_ LF (n=4), F_0_ HF (n=4), F_0_ HF-LF (n=4) male C57BL/6N mice performed using the indicated antibodies detecting mitochondrial and DICER1 proteins. HSP90 served as loading control. **(m)** Densitometric quantification of (**l)** representing n=4 biological replicates. **(n)** Illustration of proposed molecular mechanism connecting obesity and elevated *let-7* in adipocytes to post-transcriptional silencing of DICER1 and impaired oxidative phosphorylation and respiration. Two-tailed Student’s t-test **(j,k)** for each timepoint or one-way ANOVA (**d,e,h**) followed by Tukey’s multiple comparisons test were used for statistical analysis. Data are presented as mean ± standard error with individual values shown for n≤10. P-Values are indicated in the panel or represented as letters (**a**, *p*<0.05, *cel-mir-67* versus *let-7d-5p*; **b**, *p*<0.05, *cel-mir-67* versus *let-7e-5p*).

*Dicer1* 3’UTRs and exons harboured several independent *let-7* seeds of which, after mutation of putative *let-7* responsive elements (MREs), two from exonic regions became refractory to *let-7* repression, demonstrating direct negative effects of *let-7* on DICER1 translation (**Fig. 5e**). We strived to validate *let-7* targets identified in HEK293 cells adipocytes, and delivered *let-7* and control mimics to 1°eWA and confirmed that *let-7* also repressed DICER1 in white adipocytes, yet caused only modest reductions in mitochondria-/TCA-/glycolysis-associated proteins upon overexpression (**Fig. 5f-h**), prompting us to hypothesize that the chronic *let-7* induction seen in obese adipocytes might repress mitochondrial function rather via sustained DICER1 loss than by direct *let-7* effects on mitochondrial function; and that *Dicer1* inactivation might be a better proxy for mimicking the prolonged upregulation in *let-7d-5p* and *let-7e-5p* in obesity than acute overexpression of the miRNA.

To address if *Dicer1* silencing because of sustained *let-7* upregulation is sufficient to phenocopy *in vitro* the obesity-evoked impediments in mitochondrial function, we designed two independent *Dicer1-* targeting locked nucleic acid (LNA) inhibitors that repressed *Dicer1* at the mRNA and protein level, yet with varying efficiencies (**Fig. 5i**) and interrogated mitochondrial respiration using Seahorse. We found that *Dicer1* knockdown reduced oxygen consumption rates (OCR) as proxy for mitochondrial respiration (**Fig. 5j**) and extracellular acidification rates (ECAR) as proxy for glycolytic rates in 1°eWA (**Fig. 5k**) supporting our hypothesis that obesity impairs mitochondrial respiration and gene regulation, at least partially, by post-transcriptional silencing of DICER1 via *let-7* (**Fig. 5l**). *Thus, obesity-associated let-7 impairs mitochondrial oxidative respiration by silencing of DICER1 in mature adipocytes*.

## Discussion

### High transcriptional plasticity of mitochondrial dysfunction in adipose tissue

In our study we found that hepatic expression changes inflicted by HFD feeding where to a large extend irreversible, whereas adipose tissue miRNA/mRNA transcriptomes and translatomes, and particularly mitochondrial OxPhos and TCA gene sets, reverted to a state *ante*. Similar studies have investigated the effects of reversible (i.e., diet– and exercise-evoked weight gain and loss) on hepatic and adipose transcriptomes: Gonzalez-Francesa et al. observed^42^ that sustained diet-induced weight loss correlated with appreciable adipose transcriptional plasticity and mitochondrial gene expression, where short-term, exercise induced did not. Thus, durations of weight loss, and differences in control diet exposure might reflect the importance of nutritional and exposure factors during weight loss interventions on tissue responses in obesity. Despite not performing profiling of chromatic accessibility, transcriptional factor (TF) binding site enrichment analysis revealed that transcriptional activities of mitochondrial biogenesis factors like Nuclear Respiratory Factor 1 (*Nrf1*) and expression of transcriptional (co)-regulators such as Peroxisome proliferator-activated receptor gamma coactivator 1-alpha (*Ppargc1a*) and *Nrf1*^79^ were repressed in obese fat (not shown). As Brandao et al. reported ^80^ that ablation of DICER1 in adipocytes repressed *Ppargc1a*/*Nrf1*, we can conceive a scenario where reduced DICER1 (for instance due to increased *let-7d/e-5p*) drives and/or synergizes with reduced *Ppargc1a*/*Nrf1* levels to impair mitochondrial biogenesis in hypercaloric adipocytes.

### Rewiring of embryonal development by gametic miRNAs

In *C. elegans* and *D. melanogaster*, germline gene silencing by small Noncoding RNAs (sncRNA) is well-understood and similar processes are suggested for mammals ^19^. miRNAs are not only critical for sperm cell formation ^27^ and embryogenesis ^28^, but can also be transferred from sperm to zygotes and it is plausible that oocyte-sperm fusion alters zygotic miRNA pools, affect embryonic development, and instructs phenotypic traits and somatic gene expression in offspring as observed in our intergenerational paradigm. In contrast to DNA-intrinsic epigenetic modifications like DNA methylation (DNAme) and histone post-translational modifications, miRNAs diffuse between and across cellular compartments, are highly abundant in most cells, and, crucially, can be exchanged between cells through extravesicular transport ^32^. As mature sperm is transcriptionally inert, and no mechanism exists how sperm senses endocrine information and translates this into differential miRNA levels, sperm relies on ‘decoding somatic cell types’ that possess these sentinel features and are capable of instructing sperm by transfer of miRNAs and other sncRNAs like mitochondrial tRNA fragments that are exchanged between sperm and oocyte at fertilisation ^23^. Furthermore, epididymal cells, somatic epithelial cells of the extragonadal male reproductive tracts, secrete *epididysomes*, microvesicles that fuse with nascent sperm ^81^, and as we observed several miRNAs concordantly regulated in F_0_ adipose tissue and sperm, horizontal miRNA transfer from adipocytes to sperm, potentially via epididysomes is a plausible scenario that still lacks experimental proof in our paradigm. Finally, altering miRNA levels like *miR-199a-5p* in zygotes by microinjection is sufficient to reprogram embryonal metabolism ^82^, highlighting that miRNA alterations can rewire embryonal development and metabolism which are still compatible with fetal development.

## Methods

### Mice

All animal experiments were approved by the Ministry of Environment and Agriculture Denmark (Miljø-og Fødevarestyrelsen), the license no. 2018-15-0201-01544. All mice were housed under a 12-h light/ 12-h dark cycle in a temperature and humidity-controlled facility and had *ad libitum* access to diets and drinking water. 4-week-old C57BL/6J male mice (Janvier Labs) were fed with a chow diet (NIH-31, Zeigler Brothers Inc., 8% calories from fat) for 2 weeks of acclimatization and then were fed with either a low-fat diet (LF, D12450J, 10% kcal from fat, Research Diet) or a high-fat diet (HF, D12492, 60% kcal from fat, Research Diet) for 9 weeks. After 9 weeks of feeding, half of HF-fed mice were randomly assigned to the diet regression group which was given LFD. LFD is carefully chosen to match food source (animal vs plant) and micronutrient composition of HFD.

### Mouse husbandry

All male virgin mice were maintained on the assigned diet until 21 weeks of age and mated with 10-week-old C57BL/6J female virgin mice (Janvier Labs). 8-week-old C57BL/6J female mice were fed with a chow diet for 2 weeks of acclimatization and then were fed with LF during breeding and lactation. Mating was set up for 5 days in a female’s cage and vaginal plugs were checked every morning. All mice, including the F_0_ HF group, were maintained on LF during breeding to remove confounding effects from maternal diets. Males were sent from breeding cages to the original cage to be fed with the assigned diet for additional 2 weeks and then sacrificed. The offspring from all groups of founders consumed only LF. Representative male offspring from six litters per group were included in all analyses.

### Intraperitoneal glucose and insulin tolerance tests

Mice were fasted for 6 h for glucose-tolerance tests. Fasting glucose levels were measured and then mice were intraperitoneally injected with 2 g/kg body weight of 20 % glucose solution (B. Braun Medical A/S). For intraperitoneal insulin-tolerance tests, random-fed glucose levels were measured, and mice were intraperitoneally injected with 0.75 IU/kg body weight of 100IU stock insulin (Novo Nordisk). Blood glucose levels were measured at 15, 30, 60, 90, and 120 min after injecting glucose or insulin using a CONTOUR®NEXT glucometer (Bayer). For ipGTT/ipITT, we omitted mice with strongly delayed rises in glucose and no discernible drop in blood glucose, respectively, from analyses as this was the consequence of mal injection outside the peritoneum.

### Tissue collection

Mice were sacrificed by carbon dioxide euthanasia. Livers, eWAT, iWAT and BAT were weighed, snap-frozen in liquid nitrogen, and stored in –80 °C. Pieces from liver and eWAT were fixed in 4% paraformaldehyde (PFA) for histology analysis. For serum collection, blood collected by the cardiac puncture was centrifuged at 2,000 g for 15 min. The supernatant was collected and stored in –80 °C.

### Sperm collection

Mature sperm cells were collected from the cauda epididymis. Briefly, sperm cells are allowed to swim out from the cauda epididymis and incubate in PBS at 37 °C for 15 min. Supernatant containing sperms was passed through a 40-µm cell strainer and then incubated in a somatic cell lysis buffer (0.5 % Triton-X and 0.1 % SDS) for 40 min on ice. Sperms were centrifuged at 600 g for 5 min and the pellets were washed with PBS and pelted again at 600 g for 5 min. The supernatant was discarded, and the pellets were snap-frozen in liquid nitrogen and stored at –80 °C.

### Zygotic miRNA microinjection and embryo implantation

Ovulation of prepubescent (4-week-old) C57BL/6J females was induced by intraperitoneal (IP) injection of PMSG, 5 IU/female, followed by IP injection of hCG, 5IU/female, 47h later. After the second injection, females were set in breeding with C57BL/6J stud males. Next morning, they were monitored for formation of a vaginal plug (indicating mating has occurred during the night). Females were euthanized and oviducts were dissected to harvest the cumuli containing zygotes and unfertilized oocytes. Cumuli were disaggregated by incubation in a 10mg/ml solution of Hyaluronidase (Sigma-Aldrich ref. H4272) in M2 medium (Sigma-Aldrich ref. M7167) at RT for 10 minutes. Zygotes were sorted out, assessed by the presence of the second polar body and cultured in KSOM medium (Merck Millipore ref. MR-106-D). Oocytes were discarded. Each miRNA was diluted at 10 ng/ul in microinjection buffer (5 mM Tris, 0.1 mM EDTA, pH 7.4) as stock solution and kept at –80 °C. The day of microinjection, 1,5 ng/ul solutions were prepared from the stock, and injected in the pronuclei of the zygotes following standard microinjection procedures. miRIDIAN mimics were ordered from Dharmacon: Inert miRNA control is based on *C.elegans cel-mir-67* which is not expressed in mouse or human cells (5‘-UCACAACCUCCUAGAAAGAGUAGA-3‘, cat. no. CN-001000-01-05), *mmu-let-7d-5p* (cat. no. C-310486-07-0002) and *mmu-let-7e-5p* (cat. no. C-310507-07-0005). Injected zygotes were cultured in KSOM until the microinjection session was completed and then cryopreserved by the “Slow Freezing” Method (Manipulating the Mouse Embryo, a Laboratory Manual, fourth edition, Chapter 16, Richard Behringer et al. Cold Spring Harbor Laboratory Press, 2014) and transported to SDU at –150 °C for embryo transfer and recovery.

The frozen microinjected embryos were thawed at SDU and cultured in KSOM (Merck, MR-121-D) for 40 minutes (Manipulating the Mouse Embryo, a Laboratory Manual, fourth edition, Chapter 16, Richard Behringer et al. Cold Spring Harbor Laboratory Press, 2014). Viable embryos with intact zona pellucida were selected for further transfer. These embryos were then washed in 10 drops of M2 and surgically transferred to the oviducts of 0.5 dpc pseudo pregnant females. Pseudopregnancy was induced in the females by mating them with a vasectomized male. Prior to implantation, the pseudo pregnant females were anesthetized with ketamine (100 mg/kg) and xylazine (10 mg/kg) through intraperitoneal injection, and the surgical embryo transfer procedure was performed following the protocol described by Behringer et al. (Manipulating the Mouse Embryo, a Laboratory Manual, fourth edition, Chapter 6, Richard Behringer et al. Cold Spring Harbor Laboratory Press, 2014). Postoperative analgesic treatment with carprofen (5 mg/kg) was administered subcutaneously for 48 hours.

### 8-cell-morula blastomere disaggregation

miRNAs were injected in the embryo at zygote stage as described above. After injection, embryos were incubated in KSOM (KSOM-Embryomax advanced embryo medium, Merck Millipore #MD-101-D) for 48-56 hours at 37 °C, in 5% C0_2_. The culture medium was refreshed daily. In the afternoon of the third day of culture, when the embryos had reached the 8-cell morula stage, they were incubated at RT in Tyrod’s solution (Sigma-Aldrich T1788) for up to 30 seconds until the zona pellucida was completely dissolved. Afterwards the naked morulae were washed through three drops of M2 medium (Sigma-Aldrich M7167) and finally incubated in KSOM without Ca^2+^ and Mg^2+^ at RT for 20 minutes. Then, embryos were moved to a new drop of fresh M2 medium and there, pipetted up and down repeatedly to facilitate the blastomere disaggregation. When single blastomeres were obtained, they were individually transferred to single wells of a 96-well single-cell-sequencing plate from VWR (Cat. No. 82006-636) with 2 ul of RNAse inhibitor mix (containing 0.12 ul Triton X-100 10 %, 0.1 ul RNAse inhibitor (Takara (Cat. No. 2313B, 40 U/ul), 1.78 ul MilliQ water (RNAse-free)). Plates were them frozen at –80 °C until the sequencing was performed.

### Lipidomics and Metabolomics

#### Serum Extraction

100 µL of serum was mixed with 1000 µL of a 2:1 chloroform/methanol solution and 200 µL of water. Four blank samples, each containing 100 µL of water, were processed in parallel. The mixtures were shaken at 1000 rpm for 30 minutes at 4°C, followed by centrifugation at 16,000g for 10 minutes at 4 °C to separate the phases. The aqueous phase was re-extracted with 350 µL of an 86:14:1 chloroform/methanol/water solution. The organic phase from this re-extraction was combined with the corresponding organic phase from the initial extraction. Aqueous fraction was dried using SpeedVac, while the organic phase was dried under a stream of nitrogen. For subsequent analysis, the dried aqueous fractions were reconstituted in 30 µL of 1 % formic acid for metabolomics. The organic phase was reconstituted in 30 µL of lipidomic solvent A (5:1:4 isopropanol/methanol/water with 5 mM ammonium acetate and 0.1 % acetic acid). A pooled quality control sample was prepared for each analytical setup by combining 5 µL from each sample, excluding blanks.

### Lipidomic analysis

For **lipidomic** analysis, 2.5 µL of each sample was injected into an Agilent 1290 Infinity UPLC system (Agilent Technologies, Santa Clara, CA, USA) equipped with a Zorbax Eclipse Plus C18 guard column (2.1 × 5 mm, 1.8 µm) and an analytical column (2.1 × 150 mm, 1.8 µm), maintained at 50 °C. Analytes were eluted at a flow rate of 400 µL/min using eluent A (water with 5 mM ammonium acetate and 0.1 % acetic acid) and eluent B (99:1 isopropanol/water with 5 mM ammonium acetate and 0.1 % acetic acid). The gradient was as follows: 0 % B from 0 to 1 min, 0-25 % B from 1 to 1.5 min, 25-95 % B from 1.5 to 12 min, 95 % B from 12 to 14 min, and 95-0 % B from 14 to 15 min, followed by a 3-minute equilibration. For **metabolomics** analysis, 1.5 µL of each sample was injected into the same UPLC-MS system, but with a ZORBAX Eclipse Plus C18 guard column (2.1 × 50 mm, 1.8 µm) and an analytical column (2.1 × 150 mm, 1.8 µm) maintained at 40°C. Eluent A was 0.1 % formic acid, and eluent B was 0.1 % formic acid in acetonitrile. The gradient was as follows: 3 % B from 0 to 1.5 min, 3-40 % B from 1.5 to 4.5 min, 40-95 % B from 4.5 to 7.5 min, 9 5% B from 7.5 to 10.1 min, and 95-3% B from 10.1 to 10.5 min, followed by a 3.5-minute equilibration. Both analyses were performed on a 6530B quadrupole time-of-flight mass spectrometer (Agilent Technologies, Santa Clara, CA, USA) for mass spectrometric detection in positive and negative ion modes. MS1 mode settings included a scanning range of 50-1700 m/z for lipidomic and 10-1050 m/z for metabolomics, with 2 scans/sec. Internal calibration was performed using Hexakis (1H,1H,3H-tetrafluoropropoxy) phosphazene delivered through a second needle in the ion source via an isocratic pump running at 20 µL/min. Iterative data-dependent MS2 mode was used for fragmentation in both analyses with the following settings: collision energy at 40 V (lipidomics) and 20-40 V (metabolomics), precursor threshold at 5000 counts, and active exclusion after 2 spectra, with re-inclusion after 0.5 min.

### Data Processing and Normalization

All data was converted. mzML format using ProteoWizard^83^. Lipidomics data was annotated in MS-Dial (v. 4.9) against the incorporated lipid-blast^84^ with an MS1/MS2 mass tolerance of 0.005/0.01 Da and a minimum identification score of 70 %. The annotated compounds were exported to PCDL manager B.08.00 (Agilent Technologies) to create a database for area extraction in Profinder 10.0 (Agilent Technologies). Metabolomics data was processed in MzMine (v2.59) utilizing modules such as ADAP chromatogram builder and deconvolution. Compounds were annotated at Metabolomics Standards Initiative (MSI) levels 3 using the libraries of National Institute of Standards and Technology 2017 (NIST17) and MassBank of North America (MoNa). Features with signals less than 5 times those in blanks or missing in more than 20 % of QC samples were removed and signals were QC corrected using statTarget^85^.

### Data analysis

For **metabolomics** pathway enrichment analysis (KEGG pathways of Mus Musculus) comparing F_0_ LF and F_0_ HF serum (parameters ‘Scatter plot’, ‘Hypergeometric test’, ‘relative-betweenness Centrality’, ‘Use all compounds in the selected pathway library’) of differentially abundant polar metabolites (pV ≤ 0.1) were loaded into Metaboanalyst 6.0. For **lipidomics** and **metabolomics** principal component analysis and heatmap representation of indicated no. of differentially abundant lipids and metabolites were generated using ‘Statistical Feature’ (one-factor option), Sample normalization = ‘Normalization by sum‘, Data transformation = ‘Log transformation (base 10)’ and Data scaling = ‘Mean Centering‘.

### Serum insulin and leptin ELISA

Serum insulin and leptin levels were measured by mouse insulin ELISA kit (Mercodia, Uppsala, Sweden) and mouse leptin ELISA kit (Crystal Chem, IL, USA) according to manufactures’ protocol.

### Histology and staining

Liver and adipose tissues fixed with 4 % of paraformaldehyde were used for histology analyses. Formalin-fixed, paraffin-embedded tissues were cut into 3 μm sections. Sections were stained by haematoxylin and eosin (H&E) and Sirius Red by standard procedures at the Department of Pathology at Odense University Hospital, Odense, Denmark. Immunohistochemistry for CD45 was performed on Ventana BenchMark Ultra (Roche) with ready-to-use CD45 (LCA) (2B11 & PD7/26) Mouse Monoclonal Antibody.

A liver pathologist (SD) assigned steatosis, lobular inflammation, ballooning, fibrosis stage, and portal inflammation. Steatosis was graded as S0 (< 5% hepatocytes with large fat vacuoles), S1 (5-33%), S2 (>33-66%), and S3 (>66% hepatocytes with large fat vacuoles)^86^. Lobular inflammation (0-3) and hepatocellular ballooning (0-2) were assessed according to the NAS-CRN Activity Score by Kleiner et al.^86^. Also, portal inflammation was assessed, using a 4-tiered score by Kleine et al. Liver fibrosis stage was assessed according to NAS-CRN for fibrosis; F-0, no fibrosis; F-1, perisinusoidal or periportal fibrosis; F-2, perisinusoidal and portal/periportal fibrosis; F-3, bridging fibrosis; F-4, cirrhosis.

### Measurement of adipocyte volumes

The volume weighted mean volume of adipocytes was determined from the H&E sections with newCAST stereology software (Visiopharm), autodisector; measurements on systematic, randomly selected fields were performed on by at least 75 adipocyte profiles in each tissue.

### Bulk proteomics from epididymal white adipose tissue

Deep frozen fat pads (200 mg of epididymal adipose tissue) were powdered with mortar-pestle and subjected to homogenization with ceramic beads (Percellys, 5500 rpm, 2 x 10 seconds, 30 seconds break) in the buffer containing 50 mM HEPES pH 7.4, 1 % Triton X-100, 100 mM NaF, 10 mM sodium orthovanadate, 10 mM EDTA, 0.2 % SDS, 100 mM NaCl. Tissue homogenates were further processed to remove the nucleic acid traces (Diagenode SA; 10 min, cycle 30/30 seconds, at 4 °C). Floating fat and tissue debris were removed from protein extracts by centrifugation at 20000 x g. 150 mg of proteins were precipitated with 4x volume of ice-cold acetone and washed with ice-cold acetone twice. Air-dry pellets were resuspended in 8M Urea/50 mM triethylammonium bicarbonate (TAEB) buffer supplemented with a protease inhibitor cocktail (Sigma). 50 mg of the samples were incubated with 5 mM dithiothreitol at 25°C for 60 min, followed by treatment with 40 mM chloroacetamide for 30 min (room temperature, light-protected). Lys-C protease (enzyme: substrate ratio of 1:75) was added for 4 hours at 25 °C. The TAEB-diluted sample (2M Urea final) was supplemented with trypsin (1:75 ratio) followed by overnight digestion at 25 °C. Acidified peptides (1 % formic acid final) were purified on the SDR-RP (C18) multi-stop-and-go-tip (Stage Tip)^87^.

### LC MS/MS Label-Free Quantitative Proteomic Analysis from epididymal white adipose tissue

All samples were analyzed by the Proteomics Facility on a Q Exactive Plus Orbitrap mass spectrometer coupled to an EASY nLC 1000 (Thermo Scientific). Peptides were loaded with solvent A (0.1 % formic acid in water) onto an in-house packed analytical column (50 cm – 75 µm I.D., filled with 2.7 µm Poroshell EC120 C18, Agilent) and separated with 150 min gradients. Peptides were chromatographically separated at a constant flow rate of 250 nL/min using the following gradient: 4-6 % solvent B (0.1 % formic acid in 80 % acetonitrile) within 5.0 min, 6-23 % solvent B within 120.0 min, 23-54 % solvent B within 7.0 min, 54-85 % solvent B within 6.0 min, followed by washing and column equilibration. The mass spectrometer was operated in data-dependent acquisition mode. The MS1 survey scan was acquired from 300-1750 m/z at a resolution of 70,000. The top 10 most abundant peptides were isolated within a 2.1 Th window and subjected to HCD fragmentation at a normalized collision energy of 27 %. The AGC target was set to 5E5 charges, allowing a maximum injection time of 60 ms. Product ions were detected in the Orbitrap at a resolution of 17,500. Precursors were dynamically excluded for 20.0 s. All mass spectrometric raw data were processed with MaxQuant^88^ (version 1.5.3.8, using default parameters and searched against the canonical murine Uniprot reference proteome (UP589, downloaded on 28/08/2020). Label-free quantification option was enabled as was the match-between runs option between replicates. Follow-up analysis was done in Perseus^89^ 1.6.15. Hits from the decoy database, the contaminant list, and those only identified by modified peptides were removed. Afterwards, results were filtered for data completeness in replicate groups and LFQ values imputed using sigma downshift with standard settings. Finally, FDR-controlled T-tests between sample groups were performed with s0 = 0.2. Proteins were annotated on GOBP, GOCC, KEGG and Pfam terms. Afterwards, all one-way ANOVA (s= 0.2) significant candidates were Zscored and hierarchically clustered using the Euclidean distance model. Resulting clusters were merged onto the whole data set and a Fisher Exact Test calculated on cluster level with the whole dataset as background.

### RNA isolation from mouse tissue and primary adipocytes

Total RNAs from tissues, sperm, and primary adipocytes were isolated using the TRI reagent (Sigma) followed by clean-up with RPE buffer (Qiagen, Germany). The quality of RNA was validated by the Agilent RNA 6000 Nano-Kit in Agilent 2100 Bioanalyzer according to the manufacturer’s protocol. (Agilent Technologies, Waldbronn, Germany).

### RNA sequencing

*Small RNA sequencing:* (1) For sRNA-seq from F_0_ sperm, library preparation and sRNA-seq was performed at Cologne Center for Genomics, Germany. 50 ng of total RNA was used for library preparation followed by size selection (Small RNA-Seq Library Prep Kit, Lexogen). The library was sequenced in 1×50-bp single-end reads on NovaSeq6000 serie no. A00316. (2) Small RNA sequencing from F_0_ and F_1_ eWAT was carried out after quality control, 125 ng of total RNA was used for library preparation (Nextflex library kit) followed by sequencing by 1×50-bp single-end reads on Illumina Novaseq 6000. *mRNA sequencing:* mRNA sequencing was performed in-house. After quality control, 1 µg of total RNA was used for library preparation (NEBNext RNA prep kit) followed by 2×50bp paired-end sequencing on Illumina Novaseq 6000.

### RNAseq data analysis

#### Small RNA sequencing of spermatozoa

SncRNA-seq analysis of murine sperm was analyzed with Seqpac ver. 1.2.0, where sncRNA biotypes were annotated with 1 mismatch against Ensembl protein and non-coding RNA from GRCm39, miRbase, piRNA from piRBase ver. 3.0, tRNA from tRNAscan-SE, rRNA from RNAcentral and mitochondria from GRCm39:MT. Biotypes were assessed based on the following hierarchy: miRNAs, mitochondrial rRNAs, rRNAs, mitochondrial protein-coding mRNAs, mitochondrial tRNAs, tRNAs, piRNAs, nuclear protein coding mRNAs, lncRNAs, and miscRNAs. Baseline filtering of 1 read per sample and size of 15 to 75 nt long was performed. Normalization was performed with counts per million (cpm).

### Small RNA sequencing of somatic (adipose) tissue

Adapter sequences were trimmed from raw miRNA-seq reads using Cutadapt. The processed reads were mapped to the reference genome and aligned using the Bowtie algorithm within the miRDeep2 pipeline ^90^. Normalization was performed using the Trimmed Mean of M-values (TMM) method to account for sequencing depth and composition bias. Subsequent data analysis, including differential expression analysis, was conducted in R^91^ using respective Bioconductor packages^92^.

### mRNA sequencing

Sequencing reads were aligned to the mouse reference genome from Gencode (vM25)^93^ using STAR aligner (v 2.7.9a)^94^ and gene features were counted by using featureCountsv (2.0.3)^95^. The genes with low expression were filtered out using *filterByExpr* function with default settings from edgeR (v4.0.16)^96^ R package. Differential expression analyses were performed using edgeR’s *voomLmFit* ^97^unction with sample quality weights (sample.weights = TRUE) and linear model “0 + group”, whereas group represents the combination of all experimental groups of interest. The experimental conditions and linear models used for each dataset are summarized below. Genes with adjusted p-values below 0.05 were considered as differentially expressed. Gene enrichment analyses were performed by clusterProfiler (v4.10.1)^98^ R package using both over-representation analysis (ORA) and gene-set enrichment analysis (GSEA) using differentially expressed genes and log-fold-change (logFC) ranked genes (n_permutation_ = 1,000,000), respectively. Gene sets were retrieved from Gene Ontology (GO.db v3.18.0) and Reactome (reactome.db, v1.86.2) databases.

### Smart-seq2 mRNA sequencing of murine blastomeres

Custom-made Smart-seq2 protocol was implemented as before^74^. Briefly, after thawing the plate, 1 µL of dNTP mix (10 mM), 0.1 µL of oligo-dT (100 µM, /5Biosg/AAGCAGTGGTATCAACGCAGAGTAC(T)30VN-3′) and 0.9 ul nuclease-free water was added to each well and the lysis protocol was run when the samples were incubated at 72°C for 3 min in thermocycler. Samples were then put on ice for at least 1 min. Next, 5.60 µL of reverse-transcription mix containing 0.25 µL of RNase I, 2 µL of 5× first-strand buffer, 2 µL of betaine (5 M), 0.06 µL of MgCl_2_ (1 M), 0.5 µL of DTT (100 mM), 0.5 µL of SuperScript III (200 U/µL), and 0,29 ul of nuclease-free water was added to each well and reverse transcription was carried out as follows: 42 °C-90:00, 42 °C-hold, 42 °C—12:20, 10 cycles (50 °C-2:00, 42 °C-2:00), 39 °C-12:00, 70 °C-15:00, 4 °C hold. After the first 90 min of RT, 0.4 µL of 12,5 µM TSO (0.83 µM final concentration) was added to each reaction at room temperature. The plate was resealed, centrifuged for 1 min at 700 g, and placed back on the thermocycler to resume cycling from 42 °C-12:20. After reverse transcription, 15 µL of cDNA enrichment PCR reaction mix containing 12.5 µL of 2× KAPA HiFi HotStart ReadyMix, 0.25 µL of ISPCR primer (10 µM, /5Biosg/AAGCAGTGGTATCAACGCAGAGT-3′), and 2.25 µL of nuclease-free water was added to each well and cDNA enrichment PCR was performed as follows for 21 cycles: 98 °C-3:00, 7 cycles (98 °C-0:20, 60 °C-4:00, and 72 °C-6:00), 7 cycles (98 °C-0:20, 64 °C-0:30, and 72 °C-6:00), and 7 cycles (98 °C-0:20, 67 °C-0:30, and 72 °C-7:00), 72 °C-10:00, 4 °C-hold. The cDNA purification was performed using AMPure XP beads at a ratio of 1:1 (Beckmann). For elution, Qiagen elution buffer (EB) was used. cDNA concentrations were measured using the Qubit 3.0 fluorometer according to the manufacturer’s protocol, and all cDNA samples were analyzed on Agilent Tape station using the High Sensitivity DNA assay.

Dual-indexed Illumina Nextera XT sequencing libraries were prepared using 25 % of the recommended reaction volumes of Nextera XT components. All Nextera XT libraries were prepared using 125 pg of input cDNA for tagmentation, and were subsequently enriched for 12 PCR cycles, and purified using AMPure XP beads (Beckmann) at a sample:bead ratio of 0.6:1. Library concentrations were measured using Qubit 3.0 fluorometer, and the fragments were analyzed on the Agilent Tape station using the High Sensitivity DNA assay. All libraries were individually normalized and diluted to a concentration of pooled library to 5.4 ng/µL with average size ∼515bp (16 nM). The libraries were sequenced on Illumina NextSeq500 in a single-end 75-bp format (FC-404-2005, Illumina). Single-nuclei libraries were sequenced at an average depth of 3 million reads.

### Data availability – NGS

Illumina sRNA-seq and mRNA-seq datasets from F_0_ and F_1_ liver and eWAT (2) Illumina sRNA-seq from F_0_ sperm (3) Illumina mRNA-seq from offsprings sired from of miRNA-injected zygotes (F_m_ /F_MI_) eWAT (4) Illumina mRNA-seq from 1° eWA overexpressing *cel-mir-67* (negative control), *let-7d-5p* and *let-7e-5p* mimics and (5) Smart-Seq2 single-cell mRNA-seq data from miRNA-injected 8-cell embryos are available as Gene Expression Omnibus (GEO) SuperSeries GSE280278. The data are available to the reviewers’ discretion using **url**: https://tinyurl.com/3kdey4sk (**reviewer token**: inqlkwqknhanxsn)

### Data availability – Proteomics

The mass spectrometry proteomics data have been deposited to the ProteomeXchange Consortium via the PRIDE partner repository with the dataset identifier PXD042800. The data are available to the reviewer discretion: Username, Password: QHj9fv33

### Data availability – Metabolomics & lipidomics

All metabolomic and lipidomic datasets are available at the Metabolomics Workbench^99^ data repository as project ID PR002238 (https://www.metabolomicsworkbench.org).

### Male participants, sperm samples and analysis

The intervention study was approved by the Danish National Committee on Health Research Ethics (SJ-769), the Danish Data Protection Agency (REG-048-2019) and performed at the Fertility Clinic, Zealand University Hospital, Denmark^77^. The semen samples were collected from obese patients (≥ 30 kg/m_2_) via masturbation following two days of abstinence. The sperm analyses were performed in the IVF laboratory in an authorized tissue establishment. The other semen samples were diluted with MHM-C (Fujifilm) and frozen immediately for further analysis. The semen samples were collected via masturbation following two days of abstinence, and the samples were analyzed a maximum of two to three hours after production. After liquefaction, ejaculate volume and sperm concentration (total, motile and progressive) were assessed using a microscope (Leica DM750). Sperm concentration was determined with a CellVision counting chamber (CellVision Technology, Heerhugowaard, the Netherlands). Total sperm count was calculated as the product between ejaculate volume and sperm concentration. The sperm analyses were performed in the IVF laboratory in an authorized tissue establishment.

### Study design and approvals

This prospective intervention study was conducted between May 2020 and April 2022 at the Fertility Clinic, Zealand University Hospital, Denmark. The study was approved by the Danish National Committee on Health Research Ethics (SJ-769), the Danish Data Protection Agency (REG-048-2019) and registered on the Clinical Trial website (www.ClinicalTrials.gov identifier NCT04721938). All participants provided written informed consent.

### Primary adipocyte culture and transfection

Epididymal white adipose tissues (eWAT) from 6-to 8-week-old male C57BL/6J mice were removed, minced, and digested with pre-warmed collagenase media (500 U/mL collagenase type 2 (LS004183, Worthington) and 3.75U/mL DNase I (10104159001, Sigma). Isolated cells were seeded in 6-well plates and grown in DMEM/Ham’s F12 medium (21331, Vendor: Gibco), supplemented with 0.1 % Biotin/D-Pantothenate (33 mM/17 mM), 1 % penicillin-streptomycin, and 20 % FBS. Until reaching 80 % confluency, primary cells were induced by induction medium with 1 µM rosiglitazone, 850 nM insulin, 1 µM dexamethasone, 250 µM 3-isobutyl-1-methyl-xanthin (IBMX) and 10 % FBS. Subsequently, the medium was changed every other day by a differentiation medium containing 1 µM rosiglitazone, 850 nM insulin and 10 % FBS. 100 nM of Negative Control and *mmu-let-7-d/e-5p* mimics or LNA oligos targeting *Dicer1* were reversely transfected using Lipofectamine RNAiMAX (Thermo Fisher Scientific) at day 4 after induction. Completely differentiated cells were harvested at 7 days after induction for further analysis.

### Plasmid design and dual luciferase assay

For the generation of reporter constructs, the specific region of murine *Ago2*, *Dicer-1*, *Hif1an*, *Hk2*, *Cpt2*, *Crat*, *Pcx*, *Idh2*, *Sdha*, *Sdhaf3*, *Pdk2*, *Slc25a22* and *Slc25a24* containing either wildtype (wt) or mutated (mut) *let-7* binding sites was cloned behind the translation stop codon of the firefly-luciferase gene in pmirGLO Dual-Luciferase vector using custom DNA oligos provided by IDT. Renilla luciferase coding in the same plasmid was used as a control reporter for normalization purposes. To report the miRNA activity in mammalian cells, HEK 293 cells were reversely transfected in 96-well plates with 60 ng DNA using the TransIT transfection reagent (mirusbio) for 3 days and forwardly transfected with *let-7d/e-5p* mimics for 2 days. HEK-293 cells were cultured in DMEM medium (21013, Gibco) supplemented with 5.6 mM L-glutamine, 1 % penicillin-streptomycin, and 10 % FBS. After 72-h vector transfection and 48-h miRNA mimics transfection, dual luciferase reporter assays (Promega) were performed. The relative activities of firefly luciferase indicating the interaction between miRNA and specific region of targeted mRNA were normalized by the activities of Renilla luciferase.

### TaqMan assay and general qPCR

To quantify the levels of mature miRNAs, TaqMan MicroRNA Reverse Transcription Kit (4366596, Thermo Fisher), TaqMan MicroRNA Assays and TaqMan Universal Master Mix II (4440042, Thermo Fisher) were applied according to manufacturer’s protocols. Level of mature *hsa-miR-16* was used as internal control in human samples. The relative expression of mature miRNA was calculated using a comparative method (2-∂∂CT) according to the ABI Relative Quantification Method.

### Determination of oxygen consumption rates and measurement of glycolytic activity

Primary adipocytes from epididymal SVF (1°eWA) were seeded into Agilent Seahorse XFe96 Bioanalyzer microplates. Induction and differentiation were performed as described above. After four days of differentiation, 16,000 per well mature adipocytes were re-seeded and transfected with LNA oligos targeting *Dicer1* in DMEM/Ham’s F-12 medium plus 10 % Fetal Bovine Serum, 1 % P/S, 0.1 % Biotin and 0.1 % Pantothenic acid (Growth medium) at 37° C and 5 % CO_2_. Seahorse measurement was conducted at day 7 from differentiation. For each seahorse plate, the corresponding calibration plate was prepared 24 hours prior to experiments using 200 µl XF Seahorse Calibrant Agilent per well. Corresponding media were prepared before the experiment and consisted of Assay Medium supplemented with 25 mM Glucose, 1 mM Glutamine, 2 mM Sodium Pyruvate, set to pH = 7.4 and filtered sterile. The calibration plate possessed a cartridge having 4 pockets per well. Shortly before the measurement pocket A was filled with 20 µl 20 µM Oligomycin, pocket B with 22 µl 20 µM FCCP and pocket C with 25 µl 5 µM Antimycin A and Rotenone. One hour before measurements, plates were washed with 1x PBS and changed to 180 µl Assay Medium in a non-CO_2_ incubator at 37° C. Calibration was started using calibration plates, measuring O_2_ and pH LED Value/emission/Initial reference Delta for each well. All measurements started with measuring basal values, followed by injection of Oligomycin, FCCP, and Rotenone plus Antimycin A (MitoStressKit). Measurement parameters were: Mix 3 min, wait 0 min, measure 3 min with each reagent’s effect assessed within three consecutive measurement cycles with a total duration of 18 min per reagent injection.

### Protein extraction and western blot analysis

After being washed once by phosphate-buffered saline (PBS), the treated cells at harvest time were lysed by RIPA buffer supplemented with cOmplete and PhosSTOP and quantified by Pierce BCA Protein Assay Kit (23227, Thermo Fisher). Western blot analyses were carried out according to standard protocols with primary antibodies against COXIV (Cell Signaling Technologies #4850, 1:1,1000 dilution), Cytochrome C (Cell Signaling Technologies #4280, 1:1,1000 dilution), HSP60 (Cell Signaling Technologies #412165, 1:1,1000 dilution), PHB1 (Cell Signaling Technologies #2426, 1:1,1000 dilution), pyruvate dehydrogenase (Cell Signaling Technologies #3205, 1:1,1000 dilution), SDHA (Cell Signaling Technologies #11998, 1:1,1000 dilution), SOD1 (Cell Signaling Technologies #4266, 1:1,1000 dilution), VDAC (Cell Signaling Technologies #4661, 1:1,1000 dilution), HK2 (Cell Signaling Technologies #2897, 1:1,1000 dilution), CPT2 (Abcam Ab181114, 1:1,1000 dilution) and IDH2 (Cell Signaling Technologies #60322, 1:1,1000 dilution). Relative protein quantification was normalized by HSC70 (Santa Cruz sc-7298, 1:1000 dilution).

### Statistical analysis

Data were shown as mean ± standard error. One-way ANOVA and two-way ANOVA followed by Tukey’s multiple comparisons test were used for statistical analysis in *in vivo* experiments. Significant differences between groups were noted in figure legends respectively.

## Acknowledgements

We thank the CECAD Proteomics Core Facility for support with proteomics analyses, the Laboratory Manager, Stine Ravn, at The Zealand Fertility Clinic for handling and analysing the sperm samples. We are thankful for technical support from Ronni Nielsen and specialists from the Functional Genomics and Metabolism NGS team. We wish to thank Bente Møller and Irina Korshuniova from BRIC Single Cell Genomics Core Facility for excellent help for Smart-seq2 experiments. Smart-seq2 work was partly supported by CellX (The Danish Single Cell Examination Platform) funded by the Danish Research Agency through the Danish national research infrastructure program (5229-0009B). JWK, JHP and BDML received funding from the European Research Council Starting Grant TransGenRNA (#675014). JWK, CH, and NS received funding from Sygeforsikring Denmark, University of Southern Denmark and Danish Diabetes Academy that is funded by the Novo Nordisk Foundation (NNF17SA0031406). JWK received support from the NNF Challenge (#33444) and Bioscience and Basic Biomedicine Programs of the NNF (#28416). CH received grants from National Taiwan University (113L7494) and the National Science and Technology, Taiwan (114-2314-B-002-055-). MAM received support from Fundação de Amparo à Pesquisa do Estado de São Paulo (FAPESP) and Conselho Nacional de Desenvolvimento Científico e Tecnológico (CNPq), and Coordenação de Aperfeiçoamento de Pessoal de Nível Superior (CAPES). PK and MRF were supported by the Vetenskapsrådet Consolidator Grant 2022-03953 ‘InSync‘.

## Supplementary figure legends

**Figure S1:**
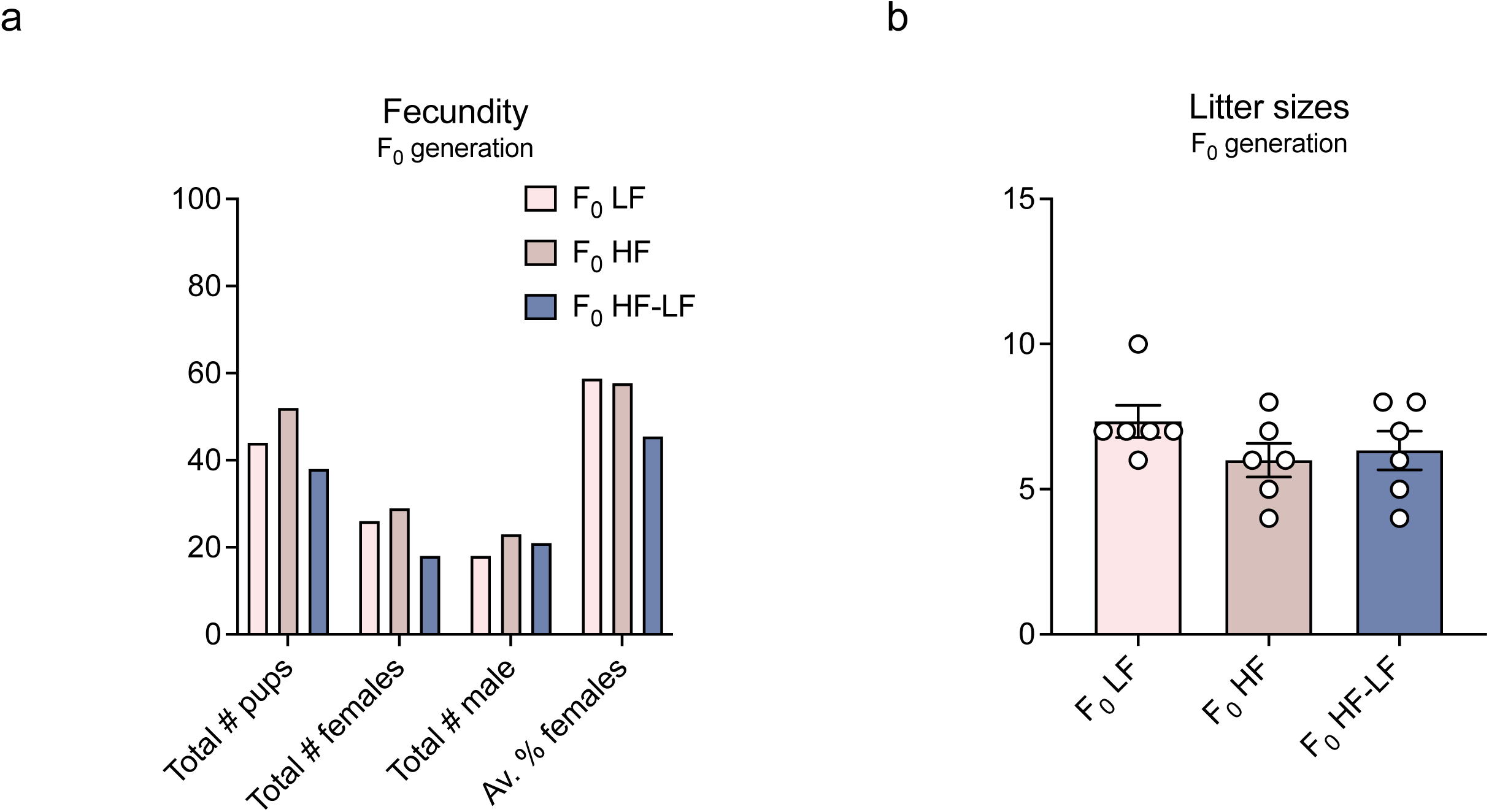
Paternal obesity and weight loss do not affect reproductive fitness and litter sizes. **(a)** Total numbers, number of male and female offsprings, sex ratios and **(b)** litter sizes of pups conceived by (F_0_ LF, n=8), (F_0_ HF, n=8) and F_1_ (F_0_ HF-LF, n=8) bucks.

**Figure S2:**
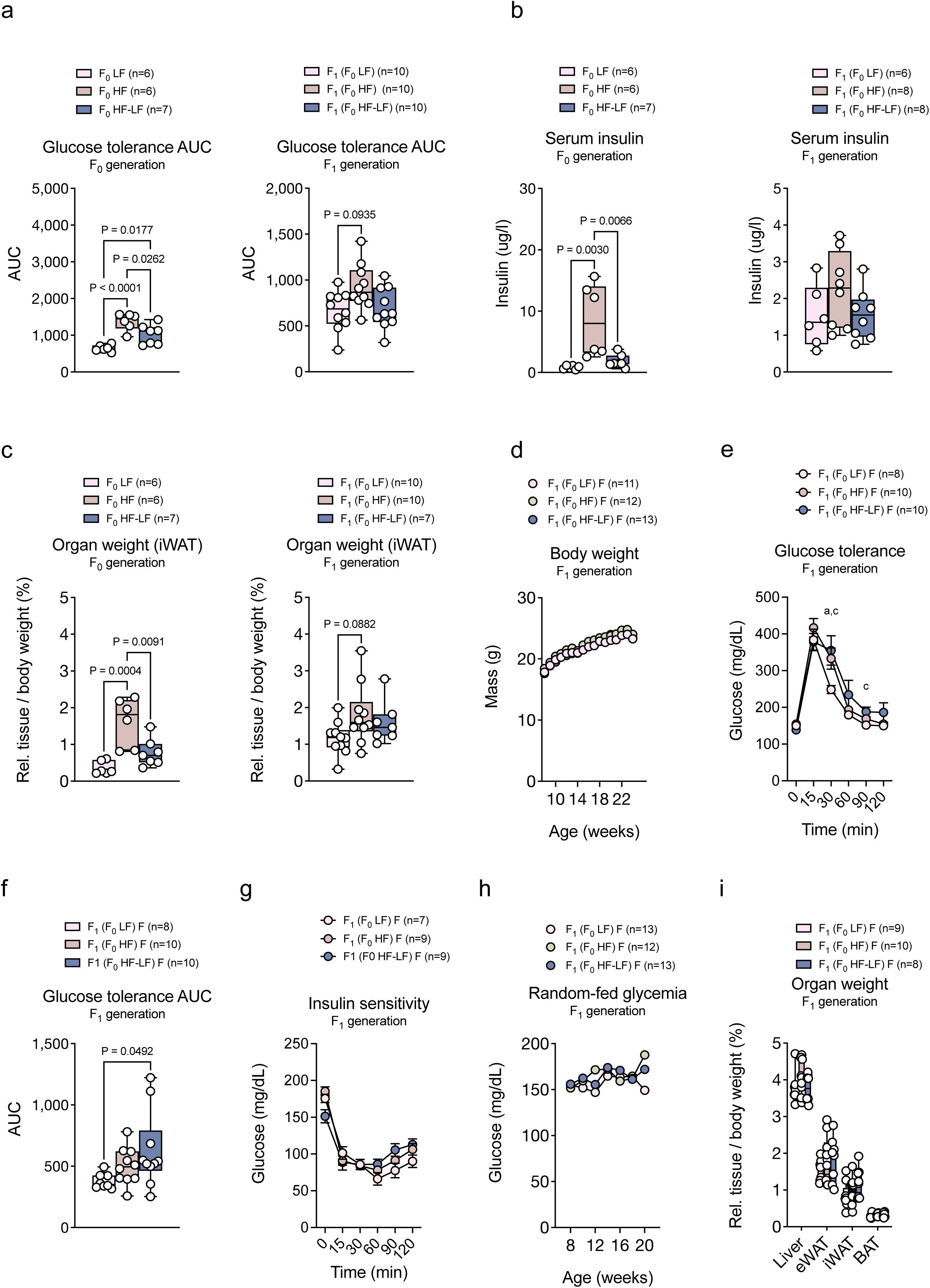
Paternal obesity causes heritable alterations in glucose metabolism in male but not in female mice. **(a)** Blood glucose area under the curve (AUC) of glucose levels during intraperitoneal glucose tolerance test in F_0_ LF (n=6), F_0_ HF (n=6) and F_0_ HF-LF (n=7, *left*) mice F_1_ (F_0_ LF, n=10), F_1_ (F_0_ HF, n=10) and F_1_ (F_0_ HF-LF, n=10, *right*) mice. **(b)** Serum insulin levels as measured by ELISA in F_0_ LF (n=6), F_0_ HF (n=6) and F_0_ HF-LF (n=7, *left*) and F_1_ (F_0_ LF, n=6), F_1_ (F_0_ HF, n=8) and F_1_ (F_0_ HF-LF, n=8) mice. **(c)** Relative inguinal white adipose tissue (iWAT) weights in F_0_ LF (n=6), F_0_ HF (n=6) and F_0_ HF-LF (n=7, *left*) and F_1_ (F_0_ LF, n=10), F_1_ (F_0_ HF, n=10) and F_1_ (F_0_ HF-LF, n=7, *right*) mice. **(d)** Body weights of F_1_ (F_0_ LF, n=11), F_1_ (F_0_ HF, n=12) and F_1_ (F_0_ HF-LF, n=13) female C57BL/6N mice. **(e)** Blood glucose during intraperitoneal glucose tolerance test in F_1_ (F_0_ LF, n=8), F_1_ (F_0_ HF, n=10) and F_1_ (F_0_ HF-LF, n=10) female C57BL/6N mice. **(f)** AUC of glucose levels during intraperitoneal glucose tolerance test in F_1_ (F_0_ LF, n=8), F_1_ (F_0_ HF, n=10) and F_1_ (F_0_ HF-LF, n=10) female C57BL/6N mice. **(g)** Blood glucose during intraperitoneal insulin tolerance test in F_1_ (F_0_ LF, n=7), F_1_ (F_0_ HF, n=9) and F_1_ (F_0_ HF-LF, n=9) female C57BL/6N mice. **(h)** Random-fed serum glucose levels in F_1_ (F_0_ LF, n=13), F_1_ (F_0_ HF, n=12) and F_1_ (F_0_ HF-LF, n=13) female C57BL/6N mice. (**i)** Relative organ weights in indicated tissue from F_1_ (F_0_ LF, n=9), F_1_ (F_0_ HF, n=10) and F_1_ (F_0_ HF-LF, n=8) female C57BL/6N mice. Two-tailed Student’s t-test (**d,e**) for individual timepoints or one-way ANOVA followed by Tukey’s multiple comparisons (**a-c,f**) were used for statistical analysis. Data are presented as mean ± standard error with individual values shown for n≤10. p-Values are indicated in the panel or represented as letters (**a**, *p*<0.05, F_0_ LF versus F_0_ HF or F_1_ (F_0_ LF) versus F_1_ (F_0_ HF); **b**, *p*<0.05, F_0_ HF versus F_0_ HF-LF or F_1_ (F_0_ HF) versus F_1_ (F_0_ HF-LF); **c**, *p*<0.05, F_0_ LF versus F_0_ HF-LF or F_1_ (F_0_ LF) versus F_1_ (F_0_ HF-LF)).

**Figure S3:**
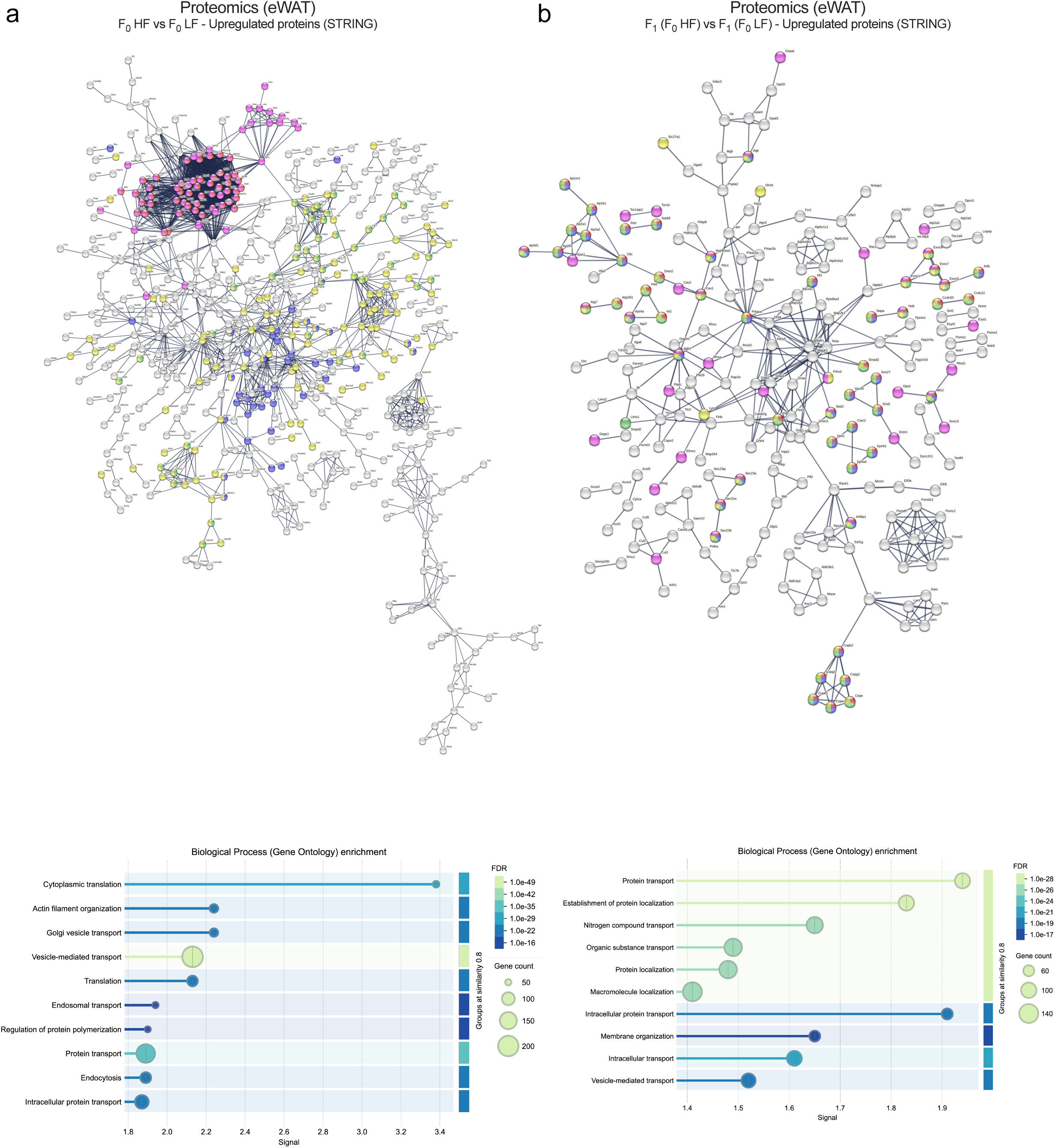
Effects of paternal obesity on F_0_ and F_1_ eWAT proteomes. (**a,b**) Visualisation of high-confidence STRING interaction networks of proteins upregulated in (**a**) F_0_ LF (n=6) vs F_0_ HF (n=6) and (**b**) F_1_ (F_0_ LF, n=6), F_1_ (F_0_ HF, n=6) eWAT (*top*) and GOBP enrichment of differentially regulated proteins (*bottom*).

**Figure S4:**
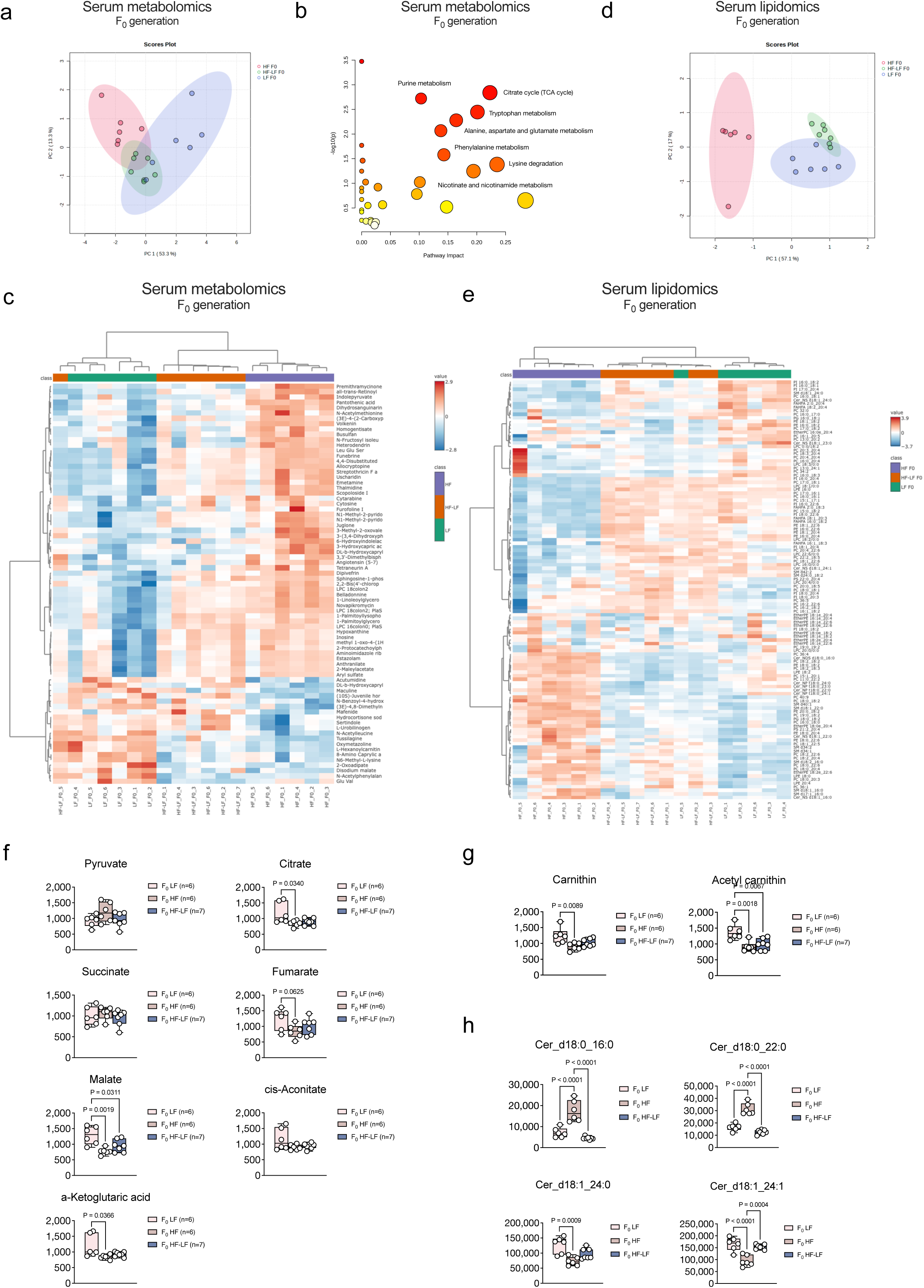
Paternal obesity elicits alterations in circulating mitochondria-associated metabolites and lipids. (**a-c**) Principal component analysis (PCA) representing PC1-2 **(a)** heatmap of the 74 most differentially regulated serum metabolites **(c)** and (**b**) GOBP term enrichment obtained by integrating differentially abundant serum metabolites and differentially expressed eWAT genes determined by RNA-seq in eWAT of F_0_ LF versus F_0_ HF male C57L/6N mice and integrated using by Metaboanalyst 6.0. (**d,e**) PCA representing PC1-2 **(d)** and heatmap of the 100 most differentially regulated serum lipids **(e)**. Metabolites and lipids were determined by ultra-performance liquid chromatography in serum of F_0_ LF compared to F_0_ HF (n=6 per group) male C57BL/6N mice. **(f-h)** Relative abundances of **(f**) mitochondrial tricarbon acid intermediates **(g)** mitochondrial Carnitine lipid carriers **(h)** mitochondria-inactivating short-chain and mitochondria-promoting long-chain ceramides in serum of F_0_ LF (n=6), F_0_ HF (n=6) and F_0_ HF-LF (n=7) mice. One-way ANOVA followed by Tukey’s multiple comparisons test were used for statistical analysis and P vales are indicated in each panel.

**Figure S5:**
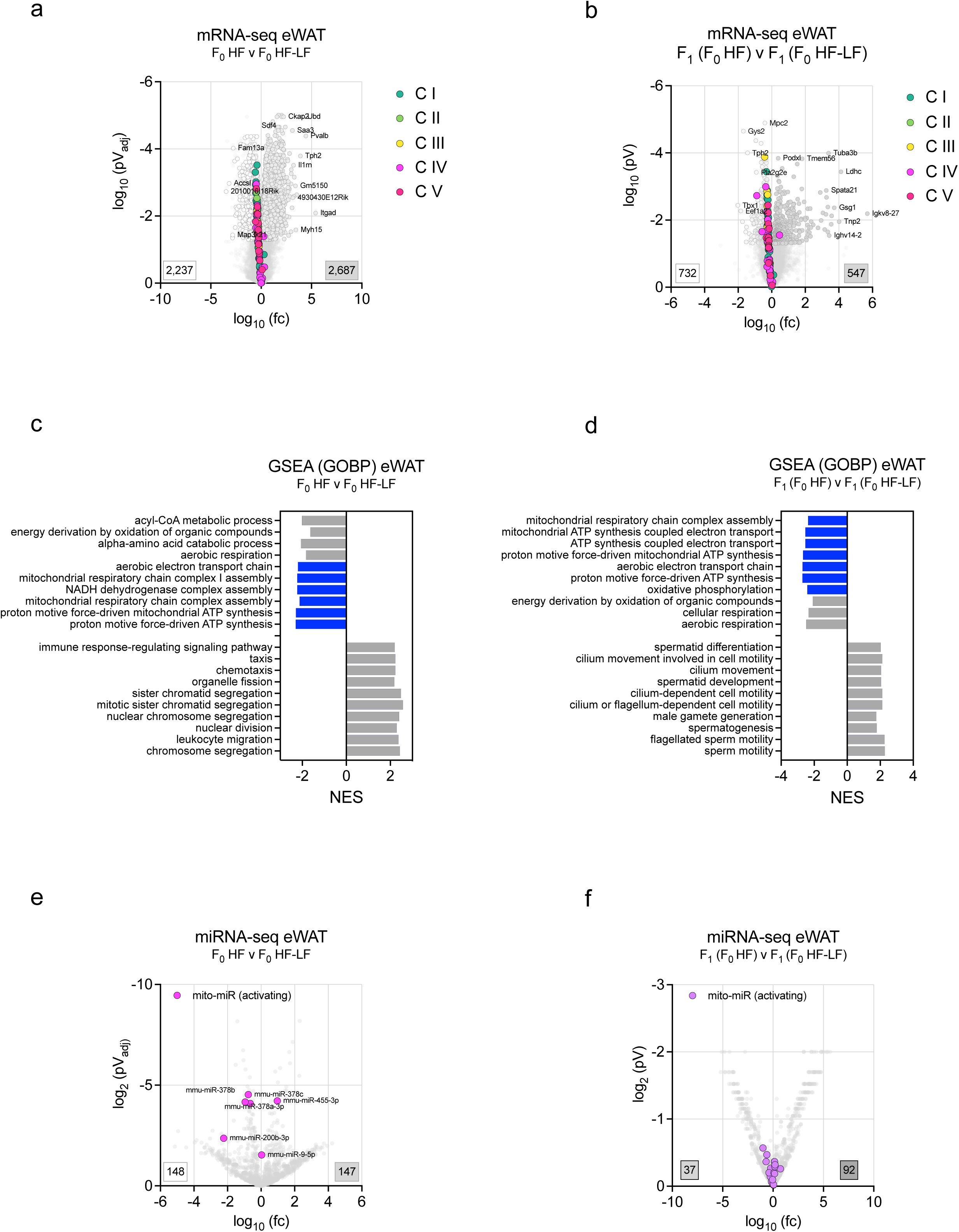
Paternal weight loss corrects obesity-associated mitochondrial dysfunction in eWAT. (**a,b**) Volcano plot of significantly up-(grey) and down-regulated (white) mRNAs in eWAT from **(a)** F_0_ HF versus F_0_ HF-LF with 2,779 up– and 1,741 down-regulated mRNAs and (**b**) F_1_ (F_0_ HF) versus F_1_ (F_0_ HF-F) with 1,118 up– and 550 down-regulated mRNAs. Plots show log10 transformed fold-changes (fc) and log10-transformed **(a)** adjusted or **(b)** non-adjusted p-Values of mRNA changes. Mitochondrial complex genes are annotated in the plot. **(c,d)** GSEA and GOBP enrichment of differentially expressed mRNAs from **(a,b).** Terms linked to mitochondrial respiration, ATP synthesis and Complex I are marked with blue bars. **(e,f)** Volcano plot of significantly regulated miRNAs in eWAT from **(e)** F_0_ HF versus F_0_ HF-F with 147 up– and 148 down-regulated miRNAs and (**f**) F_1_ (F_0_ HF) versus F_1_ (F_0_ HF-LF) with 92 up– and 37 downregulated miRNAs. Plots depict log10 transformed fc and p-Values of mRNA changes. Mitochondria-activating miRNAs are annotated in pink.

**Figure S6:**
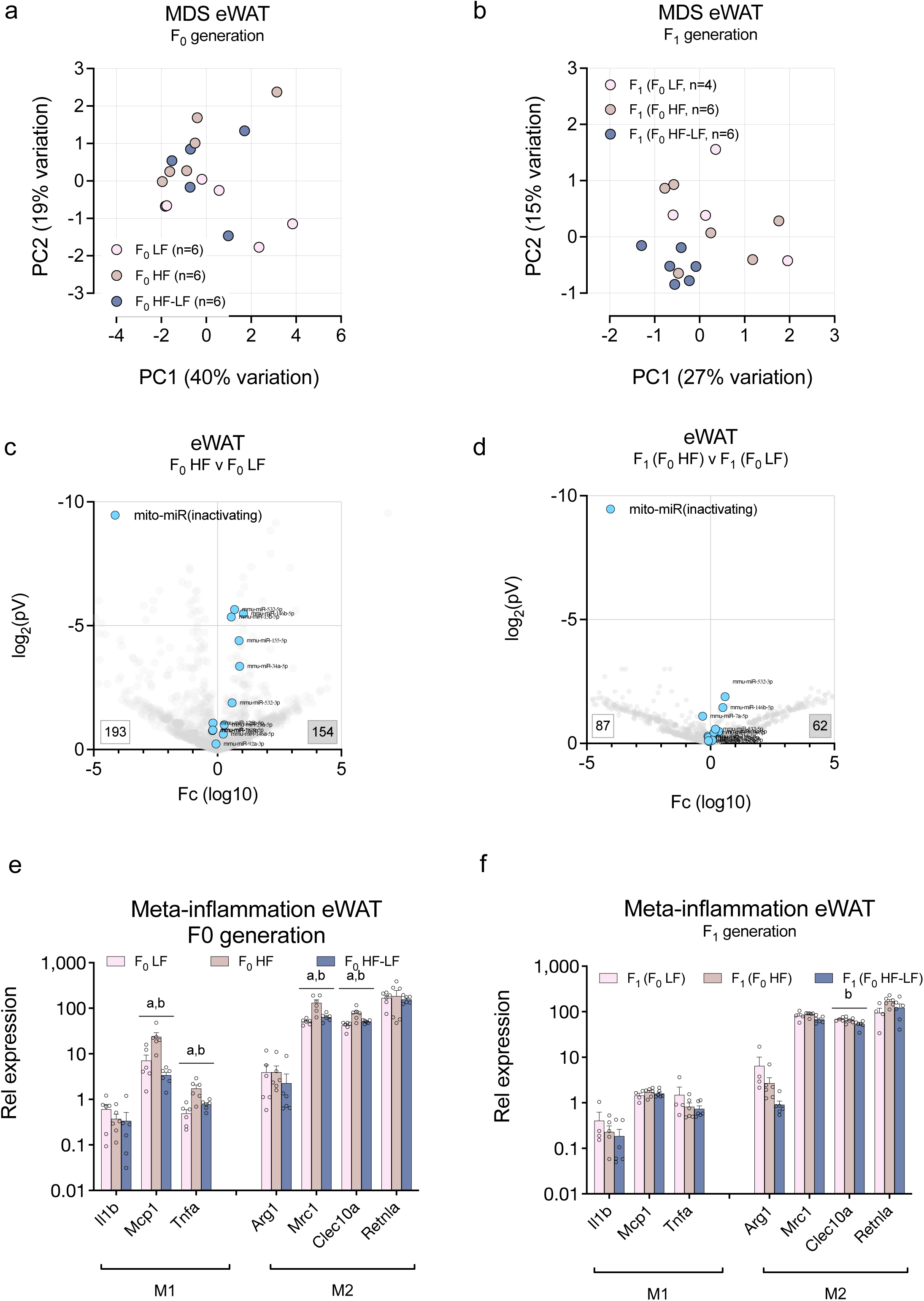
Paternal obesity does not affect F_1_ adipose miRNA expression and meta-inflammation. (**a,b)** Multidimensional scaling (MDS) plots and PC1/2 of mRNA changes in eWAT of **(a)** F_0_ LF (n=6), F_0_ HF (n=6) and F_0_ HF-LF (n=6) and (**b**) F_1_ (F_0_ HF, n=4), F_1_ (F_0_ HF, n=6) and F_1_ (F_0_ HF-LF, n=6) male C57BL/6N mice determined by mRNA-seq. **(c,d)** Volcano plot of significantly regulated miRNAs in eWAT from **(c)** F_0_ HF versus F_0_ LF with 154 up– and 193 down-regulated miRNAs and (**d**) F_1_ (F_0_ HF) versus F_1_ (F_0_ LF) with 62 up– and 87 down-regulated miRNAs. Plots depict log10 transformed fc and p-Values of mRNA changes. Mitochondria-inactivating miRNAs are annotated in light blue. **(e,f)** Expression changes reflected by normalised mRNA-seq counts from **(e)** F_0_ LF (n=6), F_0_ HF (n=6) and F_0_ HF-LF (n=6) and (**f**) F_1_ (F_0_ HF, n=4), F_1_ (F_0_ HF, n=5) and F_1_ (F_0_ HF-LF, n=6) male C57BL/6N mice. Genes represent markers of classically (M1) or alternatively (M2) activated macrophages/monocytes. One-way ANOVA followed by Tukey’s multiple comparisons (**e,f**) were used for statistical analysis. Data are presented as mean ± standard error with individual values shown for n≤10.p-Values are represented as letters (**a**, *p*<0.05, F_0_ LF versus F_0_ HF or F_1_ (F_0_ LF) versus F_1_ (F_0_ HF); **b**, *p*<0.05, F_0_ HF versus F_0_ HF-LF or F_1_ (F_0_ HF) versus F_1_ (F_0_ HF-LF); **c**, *p*<0.05, F_0_ LF versus F_0_ HF-LF or F_1_ (F_0_ LF) versus F_1_ (F_0_ HF-LF)).

**Figure S7:**
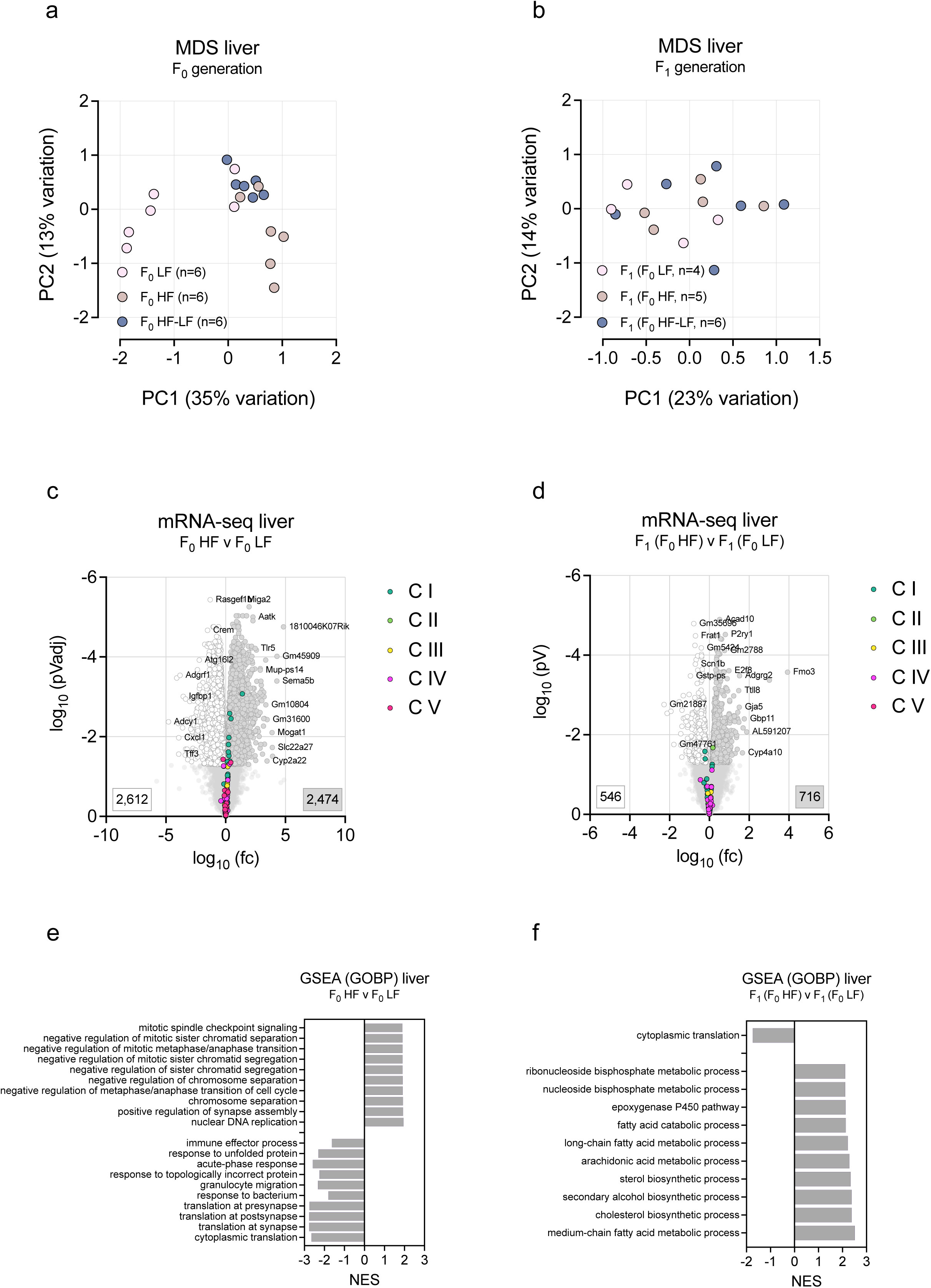
Paternal obesity does not affect mitochondrial gene expression in F_0_ and F_1_ liver. (**a,b)** MDS plots and PC1/2 of mRNA changes in liver of **(a)** F_0_ LF (n=6), F_0_ HF (n=6) and F_0_ HF-LF (n=6) and (**b**) F_1_ (F_0_ HF, n=4), F_1_ (F_0_ HF, n=5) and F_1_ (F_0_ HF-LF, n=6) male C57BL/6N mice determined by mRNA-seq. **(c,d)** Volcano plot of significantly up-(grey) and down-regulated (white) genes in liver from (**c**) F_0_ HF versus F_0_ LF, 3,312 up– and 1,597 down-regulated genes and (**d**) F_1_ (F_0_ HF) versus F_1_ (F_0_ LF), 457 upregulated and 1,041 downregulated genes. Plots depict log10 transformed fold-changes and log10-transformed **(c)** adjusted or **(d)** non-adjusted p-Values of expression changes. **(e,f)** GSEA and GOBP enrichment of biological processes expression changes shown in (**c,d**).

**Figure S8:**
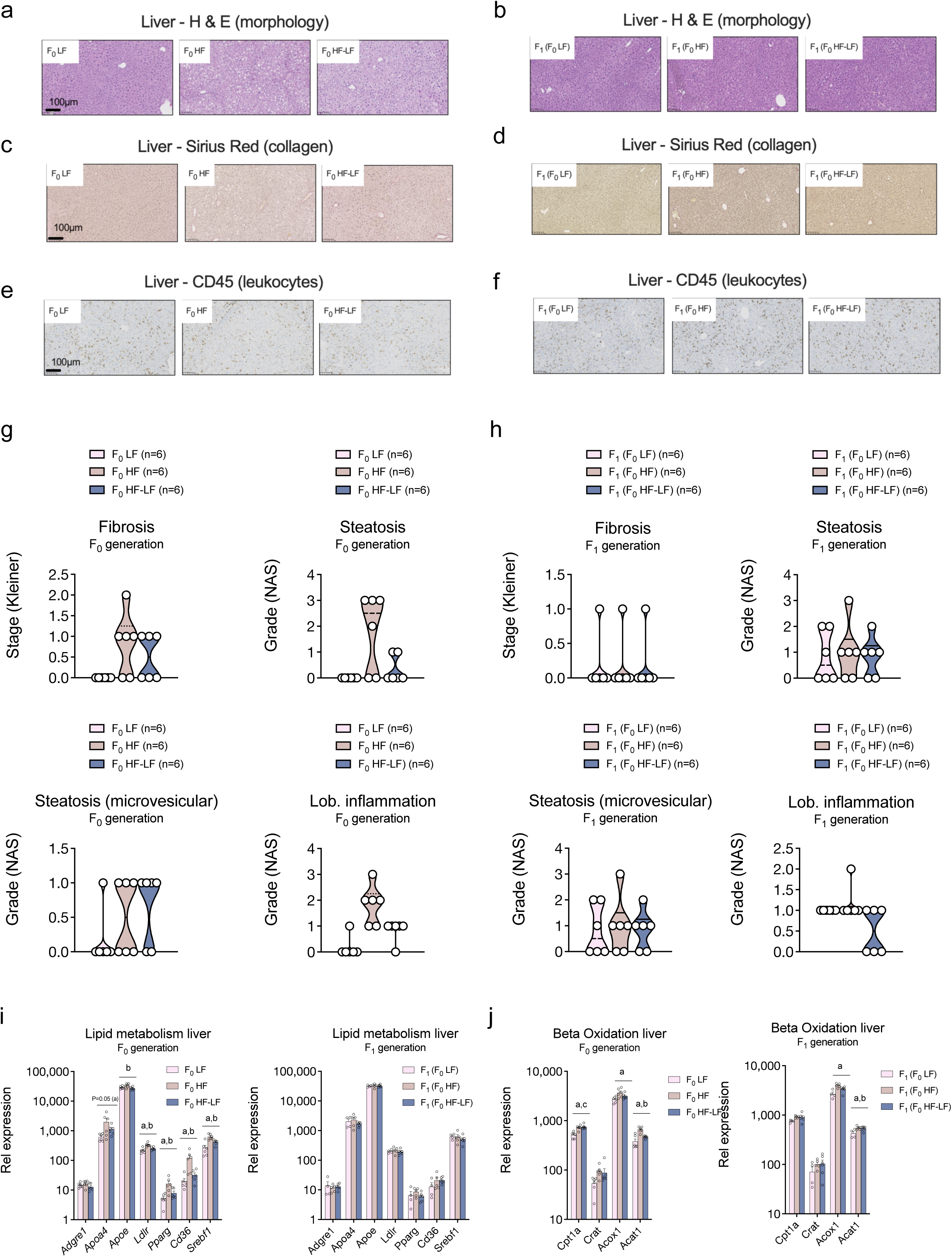
Paternal obesity and weight loss does not affect liver morphology and lipid metabolism in F_1_ progeny. (**a-f)** Representative **(a,b)** histological (haematoxylin/eosine staining cellular structures), **(c,d)** fibrosis (Sirius Red staining collagen) and **(e,f)** pan-leukocyte (anti-CD45 stains leukocytes) stained formalin-fixed paraffine embedded sections in livers of **(a,c,e)** F_0_ LF, F_0_ HF and F_0_ HF-LF and **(b,d,f)** F_1_ (F_0_ LF), F_1_ (F_0_ HF) and F_1_ (F_0_ HF-LF) male C57BL/6N mice. (**g,h**) Pathologist assessment of macrovesicular and microvesicular steatosis, lobular inflammation and NAS fibrosis scores in livers of livers of F_0_ LF, F_0_ HF and F_0_ HF-LF as well as in F_1_ (F_0_ LF), F_1_ (F_0_ HF) and F_1_ (F_0_ HF-LF, n=6 per group). **(i,j)** Expression changes reflected by normalised mRNA-seq counts from F_0_ LF (n=6), F_0_ HF (n=6) and F_0_ HF-LF (n=6, *left*) and (**f**) F_1_ (F_0_ HF, n=4), F_1_ (F_0_ HF, n=5) and F_1_ (F_0_ HF-LF, n=5, *right*) male C57BL/6N mice. Data are shown for genes in liver **(i)** lipid and lipoprotein metabolism and **(j)** mitochondrial beta-oxidation. Data in **(i,j)** are presented as mean ± standard error with individual values shown for n≤10. p-Values are shown in the panels or represented as letters (**a**, *p*<0.05, F_0_ LF versus F_0_ HF or F_1_ (F_0_ LF) versus F_1_ (F_0_ HF); **b**, *p*<0.05, F_0_ HF versus F_0_ HF-LF or F_1_ (F_0_ HF) versus F_1_ (F_0_ HF-LF); **c**, *p*<0.05, F_0_ LF versus F_0_ HF-LF or F_1_ (F_0_ LF) versus F_1_ (F_0_ HF-LF)).

**Figure S9:**
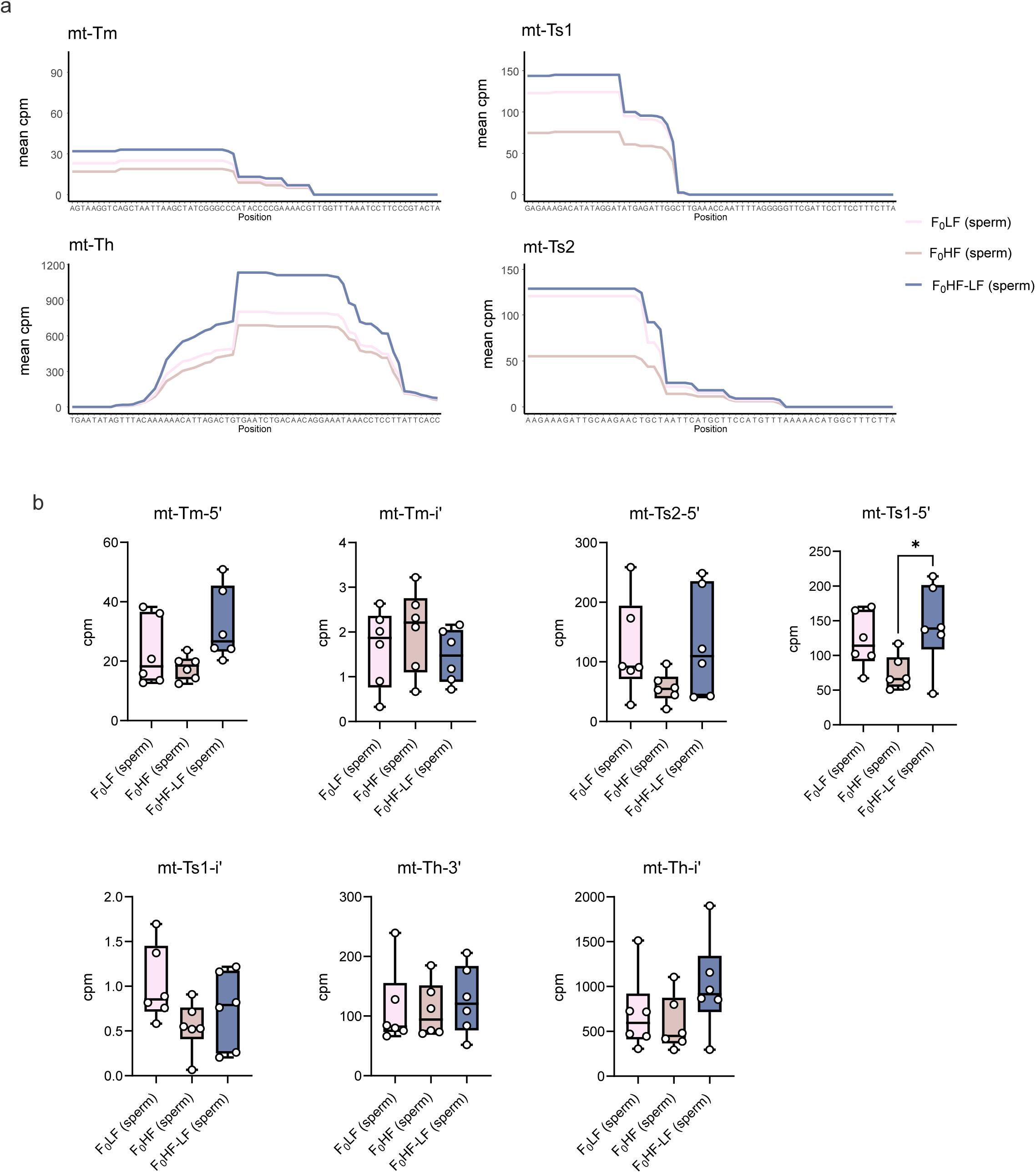
Paternal obesity and weight loss cause only modest changes in spermatozoa mitochondrial tRNA fragments. Fragments with a mean cpm above 10 shown. **(a)** Sequences mapped to mitochondrial tRNA, presented as mean cpm per diet. **(b)** Sum cpm per mitochondrial tsRNA fragment type. Each point represents one sample. (*p<0.05, One-way ANOVA with Tukey’s multiple comparisons). n=6 per diet, pink=F_0_ LF, beige= F_0_ HF, blue= F_0_ HF-LF.

**Figure S10:**
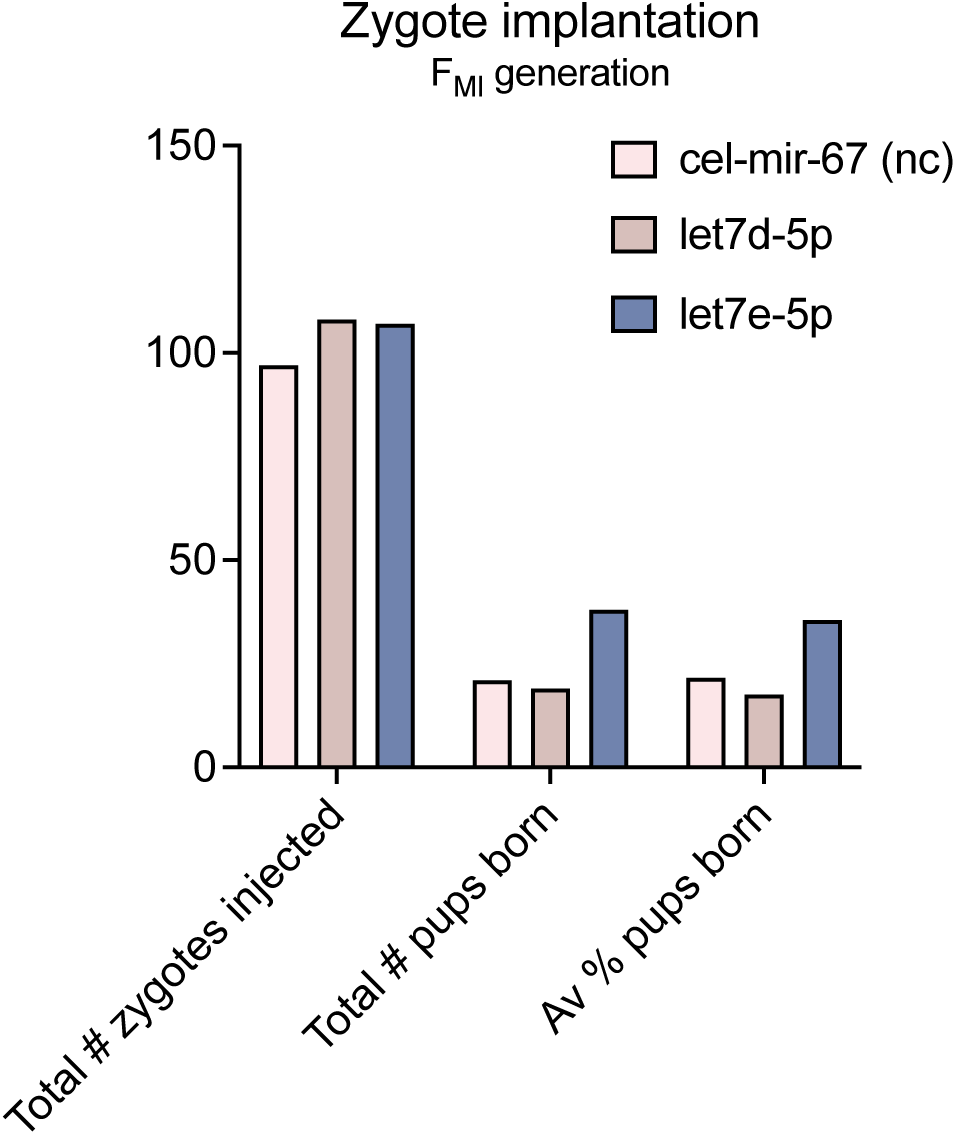
Zygotic microinjection of obesity associated *let-7* does not affect embryonic viability and implantation. Total numbers of zygotes microinjected with *cel-mir-67*, *let-7d-5p* and *let-7e-5p*, total numbers of pups born and relative survival rates of miRNA-proficient embryos.

**Figure S11:** Obesity-associated let-7 impairs adipocyte mitochondrial function via silencing of DICER1. GSEA-GOBP analysis of differentially abundant mRNAs in 8-cell blastomeres derived from *cel-mir-67* (n=22), *let-7d-5p* (n=19) and *let-7e-5p* (n=20) injected zygotes (corresponding **Fig. 4j**).

